# Neurogranin Unlocks the L-type Ca²⁺ Channel and Directs Calmodulin Delivery

**DOI:** 10.1101/2025.06.20.660829

**Authors:** Zhen Yu, Jinli Geng, Shuang Qiu, Wen Zhang, Mingjun He, Xiaodong Liu

**Author notes:** Equal contributions (Zhen Yu, Jinli Geng).

## Abstract

Neurogranin (Ng) is enriched in the postsynaptic density, with mounting evidence of its importance to neural pathophysiology. Ng has long been recognized as a dedicated calmodulin (CaM)- binding/buffering protein. This study demonstrates that Ng, through its CaM-binding IQ domain, directly binds the distal C-terminus (DCT) of the L-type Ca^2+^ (CaV1) channel, as a *de novo* mechanism regulating excitation-transcription coupling. The Ng/DCT interaction attenuates autonomous C-terminus-mediated inhibition and promotes specific CaM delivery to CaV1, cooperatively enhancing channel gating (activation and inactivation), Ca^2+^ influx and downstream neuronal signaling. Phosphorylation of Ng leads to the dissociation of the CaM C-lobe, while transiently forming the linkage with the CaM N-lobe. In contrast to neurodegenerative disease-related Ng mutants, WT Ng, within the above transient complex, enables the “targeted CaM transfer” to the proximal C-terminus of CaV1. Collective evidence from electrophysiology, binding assays, structural modeling and Ca^2+^ imaging supports a paradigm of dynamic intra- and inter-molecular interactions/facilitations underlying Ng and CaV1, both of which are representative CaM-binding proteins. This work proposes a mechanistic scheme potentially applicable to a broad scope of CaM-related physiology.

## Introduction

L-type voltage-gated Ca^2+^ (CaV1) channels, in complex with calmodulin (CaM) ^1,2^, play pivotal roles in a variety of cellular processes such as excitability, contraction, secretion and transcription ^3–5^. One of these prominent processes is the CaV1-mediated excitation-transcription/neuritogenesis coupling ^6^, where a broad spectrum of CaM- and/or growth-related signaling proteins are closely involved, e.g., the calpacitin family including neuromodulin (GAP-43) and neurogranin (Ng) enriched in the pre- and post-synapse, respectively ^7–9^. Ng is a small protein of 78 amino acids mainly incorporating the key calmodulin-binding domain (CaMBD) of the IQ motif (IQNg), widely distributed in the hippocampus and cortex of the brain ^10,11^. In line with its importance to neuronal development, Ng has been found as a critical player in neuronal plasticity, learning and memory, and neurogenerative diseases ^12,13^, all presumed by way of its role as CaM buffer dedicated to binding/releasing CaM. Beyond CaV1, CaM interacts with a broad spectrum of proteins mediating diverse signaling pathways ^5,14^. Across neurons, vascular smooth muscle cells, and cardiac myocytes, free CaM typically represents only about 1–5% of the total CaM concentration ^15–18^. More than 95% of CaM is sequestered under resting conditions. This limited availability is crucial for the fidelity and specificity of calcium-mediated signaling. How could CaM recognize its particular targets, such as CaV1 channels, instead of other CaM-binding proteins all over its close vicinity? In this work, we uncovered a direct interaction between Neurogranin (Ng) and CaV1, two prototypical CaM-binding proteins, offering new insights into the aforementioned question.

CaV1 channels are subject to CaM binding to the C-terminus (CT) of the channel, including the proximal CT (PCT) containing the CaMBD/IQ motif (e.g., IQD for CaV1.3); and the distal C-terminus (DCT), which is able to compete against CaM of its Ca^2+^ free form (apoCaM), thus switching the channel from apoCaM-bound to auto-inhibition mode ^19^. Both CaM and DCT could generate profound effects on channel functions, as manifested by CaV1.3 channels that CaM promotes ^20^ whereas DCT reduces the channel influx ^21^, with the hallmark of concurrent facilitation (CaM) or attenuation (DCT) in activation and inactivation. DCT has been suggested to resemble CaM in multiple aspects, apparently acting as a CaM-like (CaML) moiety ^6,22^. In support, DCT and apoCaM compete to bind the channel in a mutually exclusive manner. Thus far, no binding target has been reported for DCT (as CaM/CaML) except for CaV1 PCT. Meanwhile, CaM is the only protein known to directly bind Ng or IQNg, which depends on PKC (protein kinase C) phosphorylation ^9^.

Enlightened by these hints, our data unveiled that CaV1 and Ng closely resemble each other, not only in apoCaM binding but also in the key residues and mechanisms underlying DCT binding. Ng binds DCT and unlocks CaV1; meanwhile, Ng directs the CaM being unloaded from DCT-bound Ng to transfer onto CaV1 PCT, synergistically promoting CaM modulation (calmodulation). This new paradigm could potentially be expanded to other CaM-related proteins.

## Results

### Ng directly binds and promotes CaV1.3 channel activity in a phosphorylation-dependent manner

Patch-clamp data on recombinant CaV1.3 channels (with α1D as the pore-forming subunit) demonstrate that the control group exhibited CDI (Ca^2+^-dependent inactivation, quantified by *rCa*) of intermediate level (**Figure 1A**), reflecting the balance between the two basic modes of channels: CaM-bound mode with very strong CDI and DCT-bound (CaM-unbound) mode with ultraweak CDI ^19^. Overexpressing Ng could generate inhibitory effects on CaV1.3 indirectly by buffering down intracellular CaM concentrations through the high affinity of apoCaM (*Kd* = 480 nM) to the basal-state or unphosphorylated Ng ^23^. Upon Ng phosphorylation by incubating the cells with PMA, a PKC activator, the affinity for CaM would be greatly reduced (*Kd* = 19 μM), thus essentially unbinding to release CaM, altogether as the mechanism underlying the physiological role of Ng. Presumably, CDI of the PMA-treated group (CaV1.3 with phosphorylated Ng or pNg) should approach the baseline of the control group. However, significantly stronger CDI was evidenced (**Figure 1B**), which resembled that of CaM- promoted Ca^2+^ currents ^19,20^. The possibility that PMA itself would make any significant contribution to pNg effects on CaV1.3 has been excluded by nearly unaltered CDI. This surprising result suggests that a direct relationship might exist between pNg and CaV1.3, beyond the indirect effects via ambient CaM concentrations. To explore their potential interactions, Co-IP experiments were then conducted. HEK293 cells overexpressing Ng and CaV1.3 were treated with Go6983 (PKC inhibitor) or PMA, respectively. The results indicated that pNg (PMA-treated) could bind to CaV1.3, in contrast to unphosphorylated Ng (Go6983-treated) which did not yield any appreciable binding in our experimental conditions (**Figure 1C**). Collectively, Ng at the basal state (without phosphorylation) serves as a strong apoCaM buffer indirectly attenuating CaV1.3 gating; and pNg is able to directly bind to CaV1.3 presumed to be responsible for augmented CDI.

**Figure 1.**
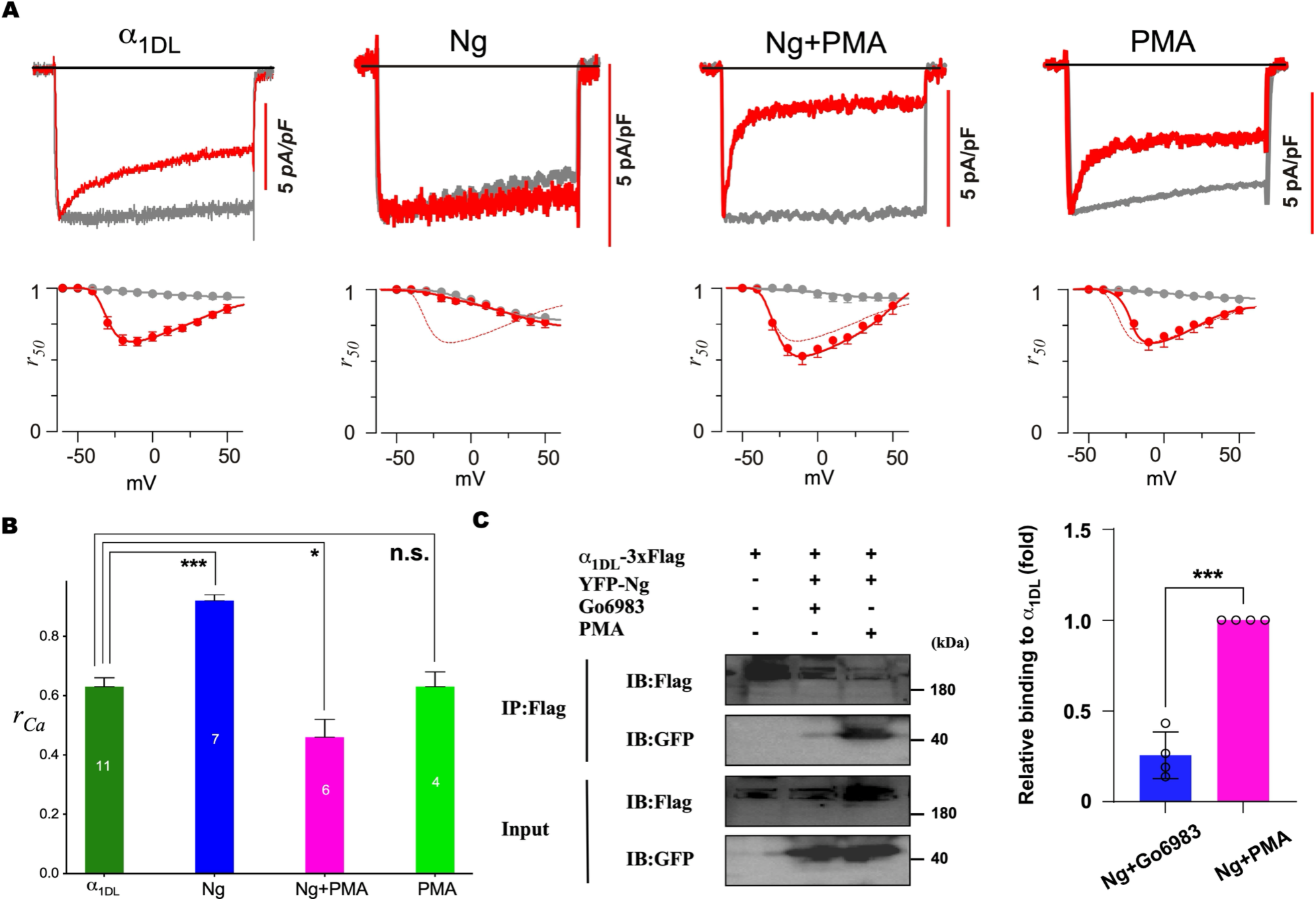
Ng modulates CaV1.3 potentially by phosphorylation-dependent binding with the channel (**A** and **B**) Ng bidirectionally modulated the inactivation (CDI) of CaV1.3 (α1DL). The Ca^2+^ currents (red) were recorded in response to a membrane potential of -10 mV for a duration of 300 ms in four groups: α1DL control, co-expression with WT Ng, WT Ng with 1 μM PMA (PKC agonist), and the PMA control (**A**, from left to right). Ba^2+^ traces recorded from the same cell were rescaled for comparison. Scale bars correspond to Ca^2+^ traces. Voltage-dependent profiles of inactivation (in Ba^2+^ and Ca^2+^) were quantified by *r50* (the ratio of the amplitudes at 50 ms versus the instantaneous peak). The ratiometric index of CDI (*rCa*, Mean ± SEM) was summarized for each group (**B**), indicated with the number of cells and the significance of difference from the control. (**C**) pNg can directly interact with α1DL. Using co-immunoprecipitation (Co-IP), the potential interactions were examined between α1DL (tagged with Flag) and Ng (with YFP), treated with either 0.5 μM Go6983 (PKC inhibitor) or 0.4 μM PMA, respectively. Anti-Flag magnetic beads were used for precipitation, followed by western blots with anti-GFP or anti-Flag antibodies. The input lane represents 10% of the total protein lysate. Representative blots are shown (left). The quantity of immunoprecipitated Ng, treated with either Go6983 or PMA, was standardized against the total amount within their respective groups (right). A minimum of three independent experiments were conducted. Statistical significance was examined (*, *p* < 0.05; **, *p* < 0.01; ***, *p* < 0.001) through one-way analysis of variance (ANOVA), followed by Bonferroni or Dunnett’s post hoc test. Values are shown as Mean ± SEM (Standard Error of the Mean).

### DCT of the channel binds pNg leading to CaV1.3 facilitation

The immediate question would be about the critical motifs from the membrane channel CaV1.3 and the signaling protein Ng, both of which are CaM-binding proteins containing the IQ domain (**Figure 2A**). Previous studies have identified the key residue of serine (S36) on Ng, which, if mutated to the negatively charged aspartic acid (S36D), would mimic the phosphorylation effects on CaM (un)binding. In contrast, the S36A mutation is immune to PKC phosphorylation. Ng_S36A and Ng_S36D were then constructed and examined for their modulatory effects on CaV1.3 channels (**Figure 2B**). Taking advantage of the two mutants that robustly represent unphosphorylated Ng and pNg, the full electrophysiological profiles were established with the indices of activation including current densities in Ca^2+^ or Ba^2+^ solutions (*JCa* or *JBa*). Consistent with the pharmacological profiles (**Figure 1**), Ng_S36A attenuated while Ng_36D enhanced CaV1.3 inactivation. In addition to the changes in inactivation (*rCa*), the amplitudes of CaV1.3 currents were decreased by Ng_S36A whereas increased by Ng_S36D. Thus, activation and inactivation were concurrently attenuated or enhanced, which, as the hallmark of DCT/apoCaM competition, strongly suggesting that DCT-mediated autoinhibition is closely involved. To systematically explore the intracellular domains of CaV1.3 for the key binding target, FRET two-hybrid binding experiments were conducted, between the donor CFP-Ng _S36D and the YFP-tagged acceptors, i.e., I-II loop, II-III loop, III-IV loop, PCT, or DCT (represented by PCRD-DCRD) (**Figures 2S1** and **2S2**). By fitting the binding curves, PCRD-DCRD exhibited the strongest affinity to Ng_S36D (equivalent *Kd* = 6.75×10^4^), among all the intracellular domains being examined (**Figure 2S3A**). In this context, it is plausible that pNg (Ng_S36D) would bind CaV1.3 at its DCT, therefore attenuating channel autoinhibition. Further Co-IP experiments clearly demonstrated that Ng_S36D was able to bind CaV1.3 and DCT, in contrast to Ng_S36A which merely exhibited any sign of interaction (**Figure 3C** and **3D**). Using FRET, Ng_S36D was confirmed to bind PCRD-DCRD with high affinity. In contrast, Ng_S36A, mostly occupied by apoCaM (due to its high affinity) under normal cell conditions of free CaM, did not bind PCRD-DCRD (**Figure 3E**). To reassure the phosphorylation-dependent effects under the conditions in **Figure 1**, the interactions between DCT and Ng were examined in HEK293 cells with PKC activator (PMA) and inhibitor (Go6983), respectively. Both Co-IP and FRET results confirmed that phosphorylated Ng (PMA-treated) could bind DCT substantially stronger than unphosphorylated Ng (Go6983- treated) (**Figure 2S3B** and **2S3C)**.

**Figure 2.**
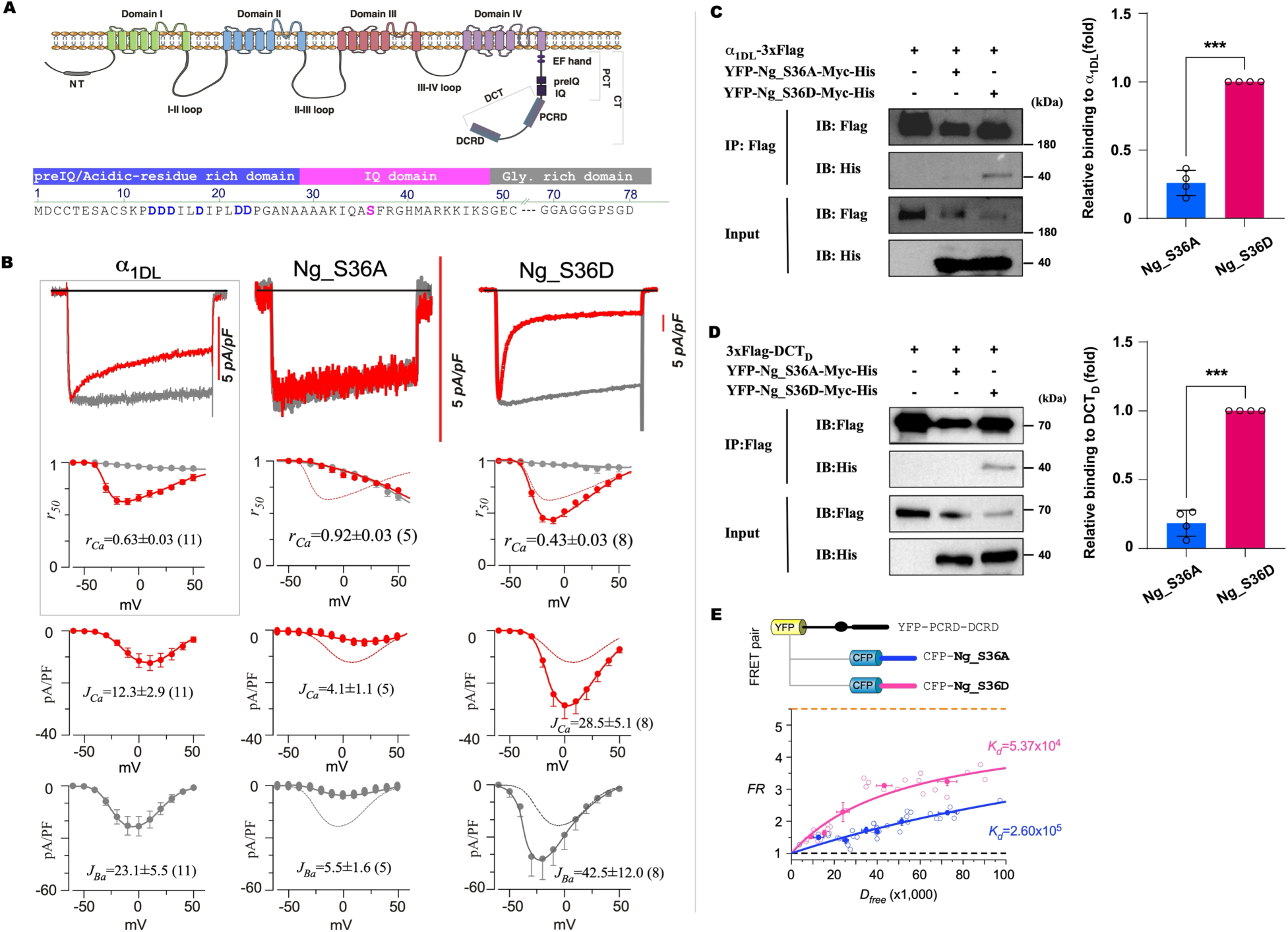
DCT mediates the direct interaction between Ng and CaV1.3 (A) Topological illustrations of α1D (upper) and Ng (lower). The four transmembrane domains (I-IV) of the α1D channel are connected by the intracellular N-terminal domain (NT), the intracellular loops (I-II, II-III and III-IV), and the C-terminal domain (CT), which includes the key domains of PCT (containing preIQ and IQ, etc.) and DCT (containing PCRD and DCRD, etc.). The scheme of Ng illustrates the major domains including preIQ/acidic-residue rich domain (blue), IQ domain (pink), glycine-rich region (GCR, gray). The key residues, including the phosphorylation site S36 (pink) and several acidic residues (D, blue) in preIQ, are highlighted. (B) The effects of the two Ng mutants on CaV1.3 currents. The traces were examined for channel inactivation for the groups of either Ng_S36A or Ng_S36D all co-expressed with CaV1.3 channels. (C) Interactions between the Ng mutants and CaV1.3. The potential interactions between α1DL (tagged with Flag) and Ng_S36A or Ng_S36D (both tagged with His) were examined by Co-IP. Representative blots (left) and the statistical summary (right). (D) Interactions between the Ng mutants and CaV1.3 DCT. The potential interactions between DCT from α1D (tagged with Flag) and Ng_S36A or Ng_S36D (both tagged with His) were examined. Representative blots (left) and the statistical summary (right). (E) FRET 2-hybrid binding assay to analyze DCT and Ng interactions. CFP-Ng_S36D and YFP- PCRD-DCRD resulted in the binding curve of *FR-Dfree* (*FR*: FRET ratio; *Dfree*: free donor concentration, shown in pink) with an apparent affinity (*Kd* = 5.37 × 10^4^), in comparison with CFP-Ng_S36A (*Kd* = 2.60 × 10^5^, blue). Each data point (solid with error bars) represents the average over five cells (smaller circles).

**Figure 3.**
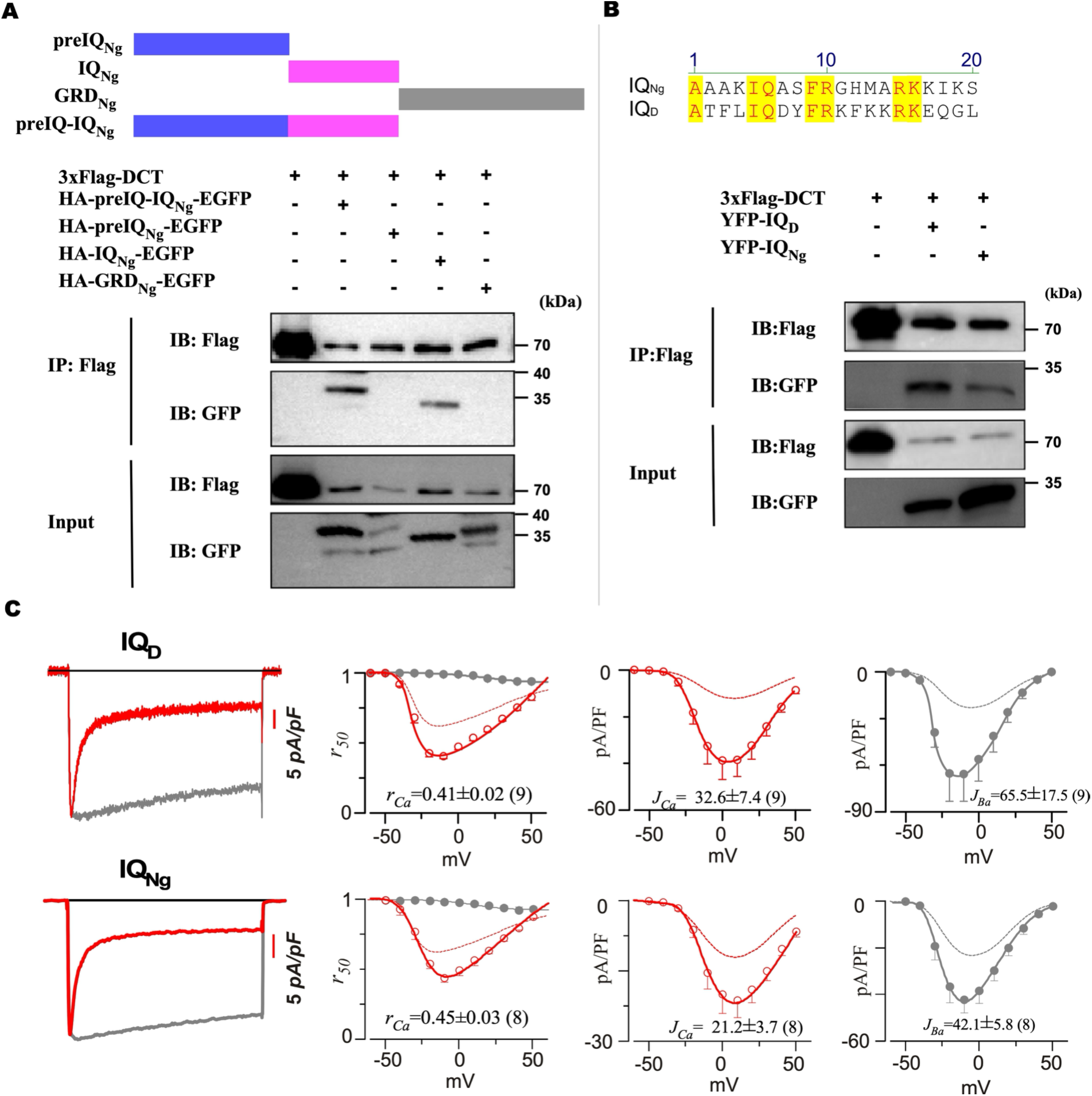
IQ motif of Ng directly binds CaV1.3 DCT (A) Ng mutants with truncations of the key domains (upper). Potential interactions between DCT (from α1D, tagged with Flag) and various peptides encoded by different domains of Ng (tagged with HA) were examined (lower). (B) Alignment of the IQ domains from Ng and CaV1.3 (upper). Co-IP results comparing YFP- IQD and YFP-IQNg for the potential interactions with DCT (tagged with Flag, lower). (C) Both IQNg and IQD promote α1DL channels. The electrophysiological profiles (from left to right), including the current traces, *rCa* (inactivation), *JCa* and *JBa* (activation) were compared among the three groups: α1DL control, α1DL overexpressed with IQNg or IQD.

### IQ motif of Ng directly binds DCT and facilitates CaV1.3

As illustrated in **Figure 2A**, the Ng protein consists of several functional domains including the preIQ domain (rich in acidic residues), the IQ domain and the glycine-rich domain (GRD). To identify the critical domain in Ng/DCT interactions, a series of HA-tagged truncation mutants were constructed, including preIQ-IQ, IQ, GRD or their combinations, and their interactions with Flag-tagged DCT were explored by Co-IP. The two constructs, preIQ-IQ and IQ, stood out as the potential targets, which were further narrowed down to the IQ motif (**Figure 3A**). It is notable that the two (apo)CaM-binding proteins, CaV1 and Ng, both have the crucial IQ domain and the negatively-charged preIQ domain, both of which are implicated in CaM binding ^24,25^. Indeed, IQNg and IQD have a high degree of sequence homology (**Figure 3B**). Moreover, both were able to bind DCT demonstrated by Co-IP results. Importantly, both IQD and IQNg significantly enhanced CDI (*rCa*) of CaV1.3, and concurrently promoted activation (*JCa* or *JBa*) (**Figure 3C**). Taken together, multiple lines of evidence strongly suggest that the IQ domain binds DCT or CaM in a mutually-exclusive manner, as the mechanism underlying pNg/CaV1.3 interaction and facilitation. Upon phosphorylation, Ng (turning into pNg) unbinds apoCaM, thereby exposing the IQNg domain to bind DCT. This interaction attenuates the autonomous DCT/IQD interaction or DCT/apoCaM competition, shifting the balance among CaV1.3 channels and resulting in a greater proportion of channels in a CaM-bound mode, characterized by strong activation and inactivation. In addition, these data also support the previously proposed hypothesis that DCT likely resembles CaM functionally, biochemically and structurally ^6^. It became imperative to investigate into the binding details of IQ/DCT at the interface between Ng and CaV1.3.

### Molecular details underlying IQ/DCT binding

First, based on the fact that IQD and IQNg demonstrate substantial similarities in coding sequences (>6 conserved residues out of 20 a.a., **Figure 4A**), DCT interactions (**Figure 3B**) and CaV1 modulations (**Figure 3C**), alanine scanning was conducted by introducing double- or single-point mutations (IQD as the template) to identify the key residue(s) critical to binding and modulation of CaV1.3. Electrophysiology was performed to pinpoint which mutant variant would effectively reduce/eliminate CaV1.3 facilitation (**Figure 4A**). Neither one of alanine substitutions on IQ, i.e., IQ/AA, I/A or Q/A, was able to cause any appreciable difference in CDI from that of WT IQ. The IQ domain is well known for its critical role in binding CaM of both apo or Ca^2+^ forms ^5^, indicating that despite the similarity between DCT and CaM, they clearly have some difference in details on their interactions with IQ. Similarly, RK related mutations did not significantly affect the strong IQ effects on CDI of CaV1.3. In contrast, double mutations of FR/AA significantly reduced the strong CDI associated with the IQ peptides. Single-point F/A and R/A mutations proved that the phenylalanine (Phe or F) at the 9^th^ position (corresponding to F37 of Ng) is the critical residue responsible for the observed CDI facilitated by IQ. Further in-depth analyses of the IQD_F/A peptide confirmed that F/A disrupted the facilitation of both inactivation and activation (the full profiles of *rCa* and *JCa* or *JBa*) by WT IQD (**Figure 4B**). Similar results were obtained from the equivalent mutant of IQNg (IQNg_F/A), consolidating the importance of the Phe residue (9^th^ in IQ or 37^th^ in Ng). Structural modeling suggested that the two binding pairs of IQD/DCT or IQNg/DCT resemble each other including the putative side-chain interactions with Phe (**Figure 4S1**). Moreover, regarding the functional effects of full-length pNg, Ng_S36D_F37A was compared with Ng_S36D, which resulted in the same conclusion: F37A disrupted pNg/CaV1.3 facilitation and presumably also their interaction. To examine the latter, FRET binding assays were conducted to measure the affinities associated with different binding pairs, for which CFP-DCRD served as the donor, and YFP-[IQD] and the FR/AA or F/A mutant as the acceptors (**Figure 4C**). Compared with YFP-[IQD] (*Kd* = 4.06x10^3^), both mutants YFP-[IQD]_F/A and YFP-[IQD]_AA exhibited weaker interactions with CFP-DCRD (*Kd* =6.81x10^3^ or *Kd* =1.03x10^4^, respectively). In the context of the full-length Ng, the double mutant CFP-NgS36D_AA and the single mutant CFP-NgS36D_F37A resulted in similar changes in binding affinities (**Figure 4D**). The binding affinities between DCT (YFP-PCRD-DCRD) and pNg mutants (CFP- NgS36D_F37A: *Kd* = 1.40x10^5^; CFP-NgS36D_AA: *Kd* = 1.60x10^5^) were significantly reduced in comparison with WT pNg (CFP-Ng_S36D, *Kd* = 6.65x10^4^).

**Figure 4.**
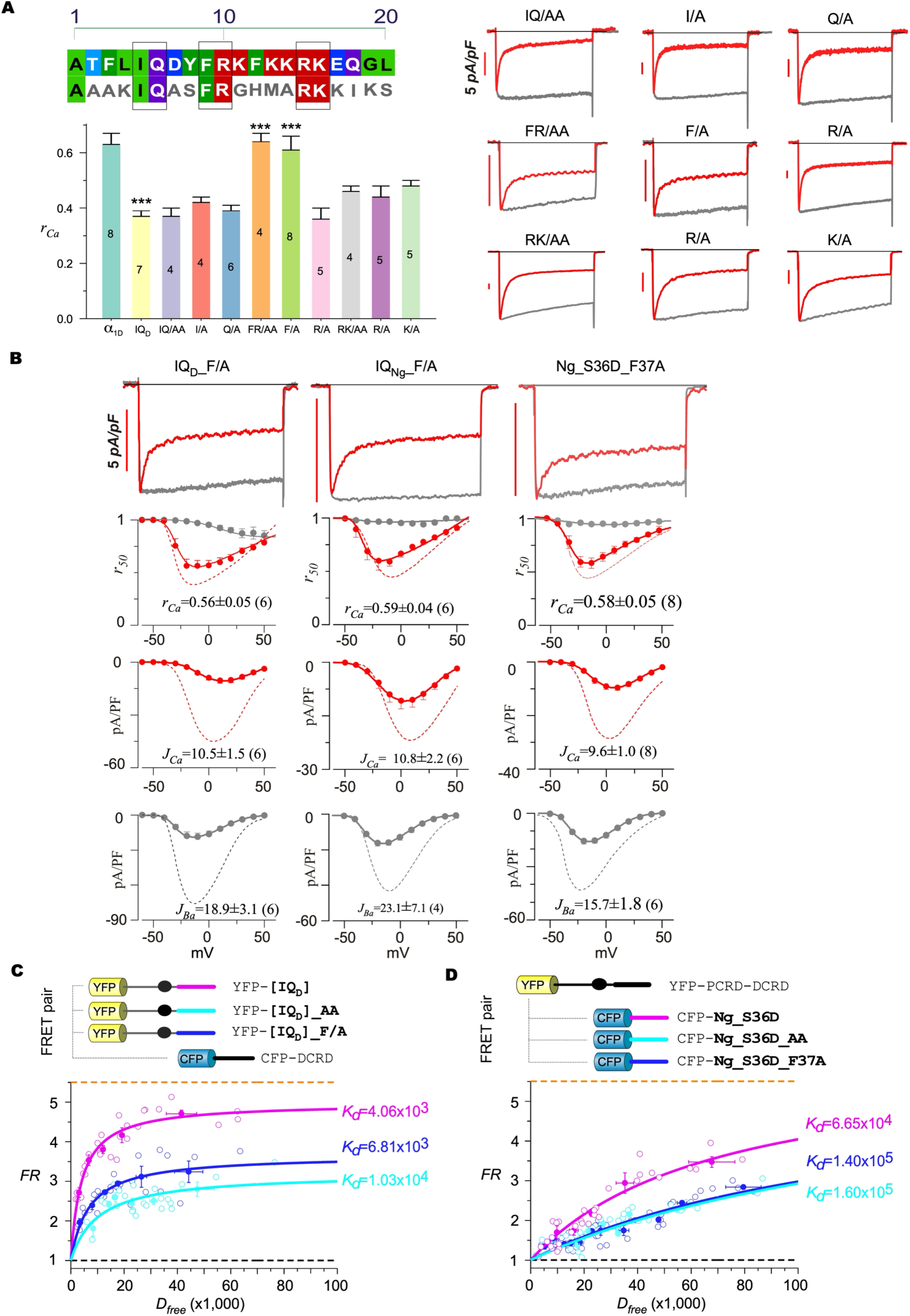
Key residues at the interface between IQ and DCT (A) Mutagenesis screening based on the key residues shared by IQNg and IQD (indicated by three box frames). The key mutations (double- or single-point) are summarized to compare the peptide effects on CDI (left) based on the current recordings of the mutants (right). (B) The electrophysiological profiles (from top to bottom), including the current traces (Ba^2+^ and Ca^2+^), *rCa* (inactivation index), and *JCa* and *JBa* (activation of Ca^2+^ and Ba^2+^ currents), for IQD_F/A, IQNg_F/A or Ng_S36D_F37A co-expressed with α1DL. (C) Interactions between IQD with key mutations and DCT examined by FRET. Binding curves were acquired between CFP-DCRDD and the two mutants of YFP-preIQ-IQD-PCRDD (i.e., YFP- [IQD]): AA or F/A. CFP-DCRDD and YFP-[IQD] resulted in the binding curve of *FR-Dfree* (pink) with apparent affinity (*Kd* = 4.06 × 10^3^), in comparison to CFP-DCRDD and YFP-[IQD] _F/A (*Kd* = 6.81 × 10^3^, blue), and CFP-DCRDD and YFP-[IQD]_AA (*Kd* = 1.03 × 10^4^, cyan). Each data point (solid with error bars) was averaged over five cells (smaller circles). (D) Key residues of IQNg for DCT/IQ interactions were examined by FRET. Binding curves were acquired between YFP-PCRD-DCRD (equivalent to YFP-DCT) and the two mutants of CFP-Ng_S36D (i.e., CFP-Ng_S36D_AA or CFP-Ng_S36D_F37A). CFP-Ng_S36D and YFP- PCRD-DCRD resulted in the binding curve of *FR-Dfree* (pink) with apparent affinity (*Kd* = 6.65 × 10^4^), in comparison to CFP-Ng_S36D_F37A and YFP-PCRD-DCRD (*Kd* = 1.40 × 10^5^, blue), and CFP-Ng_S36D_AA and YFP-PCRD-DCRD (*Kd* = 1.60 × 10^5^, cyan). Each data point (solid with error bars) represents the average over five cells (smaller circles).

### Expanding binding pairs with facilitation across the CaM-binding protein superfamily

As we demonstrated, the two CaM-binding proteins (Ng and CaV1.3) are able to bind with each other to achieve functional enhancement. Whether such mode of interaction and facilitation could be generalized to the superfamily of CaM-binding proteins is an intriguing question. We first explored the calpacitin family for the potential expansion. The IQ domain from another calpacitin protein, Nm (neuromodulin), was examined by Co-IP along with IQNg and IQD, which unveiled that all the three peptides could bind DCT of CaV1.3 (**Figure 5A**). Regarding functional effects, Nm and CaV1.3 were co-expressed in cells and then treated with PMA (**Figure 5B**). Prior to PMA treatment, Nm nearly diminished the CDI of CaV1.3 due to CaM-buffering of unphosphorylated Ng, whereas phosphorylated Nm significantly enhanced the CDI of CaV1.3. To validate this effect, Nm_S41A and Nm_S41D, representing Nm before and after phosphorylation, were also examined by electrophysiology. Compared to the control group of CaV1.3, CDI was decreased or increased by Nm_S41A and Nm_S41D respectively, which faithfully reproduced those phenotypes previously achieved from Ng mutants (Ng_S36A and Ng_S36D). It is reasonable to speculate that IQ or IQ-like domains from other CaM-binding or IQ- containing proteins may behave similarly to bind DCT of CaV1.3 and enhance channel functions, providing a new paradigm of modulation potentially adopted by a vast number of signaling proteins. On the other hand, the targeted protein of modulation could readily expand onto other channels or proteins containing EF-hands or CaM-like domains, such as other CaV1 subtypes. Employing FRET 2-hybrid assays, the binding interactions between Ng_S36D and CaV1.2-1.4 were examined (**Figure 5C**). The results indicated that all the three types of DCT peptides were able to bind Ng_S36D, with the affinities of varying levels, suggesting that the CaV1 channels should share the same principle of Ng modulation but with quantitative differences. CaV1.2 is particularly intriguing to explore further with electrophysiology (**Figure 5D**), as a subtype of widely expressed L-type CaV1 channels also subject to CMI ^6,26^. When co-transfecting IQ and CaV1.2 in HEK293 cells, similar effects of channel facilitation were evidenced from the key indices of Ca^2+^ current recordings. The promotion of CaV1.2 activation was as pronounced as 4-fold increase in current density (*JCa*). Meanwhile, IQ clearly enhanced inactivation (*rCa*) of CaV1.2, known to exhibit relatively strong inactivation even under the basal conditions. Taken together, the successful expansions from Ng/CaV1.3 onto other calpacitin and CaV1 subtypes strongly suggest that more intermolecular events pertaining to CaM/IQ or CaM/IQ-like domains await to be unveiled.

**Figure 5.**
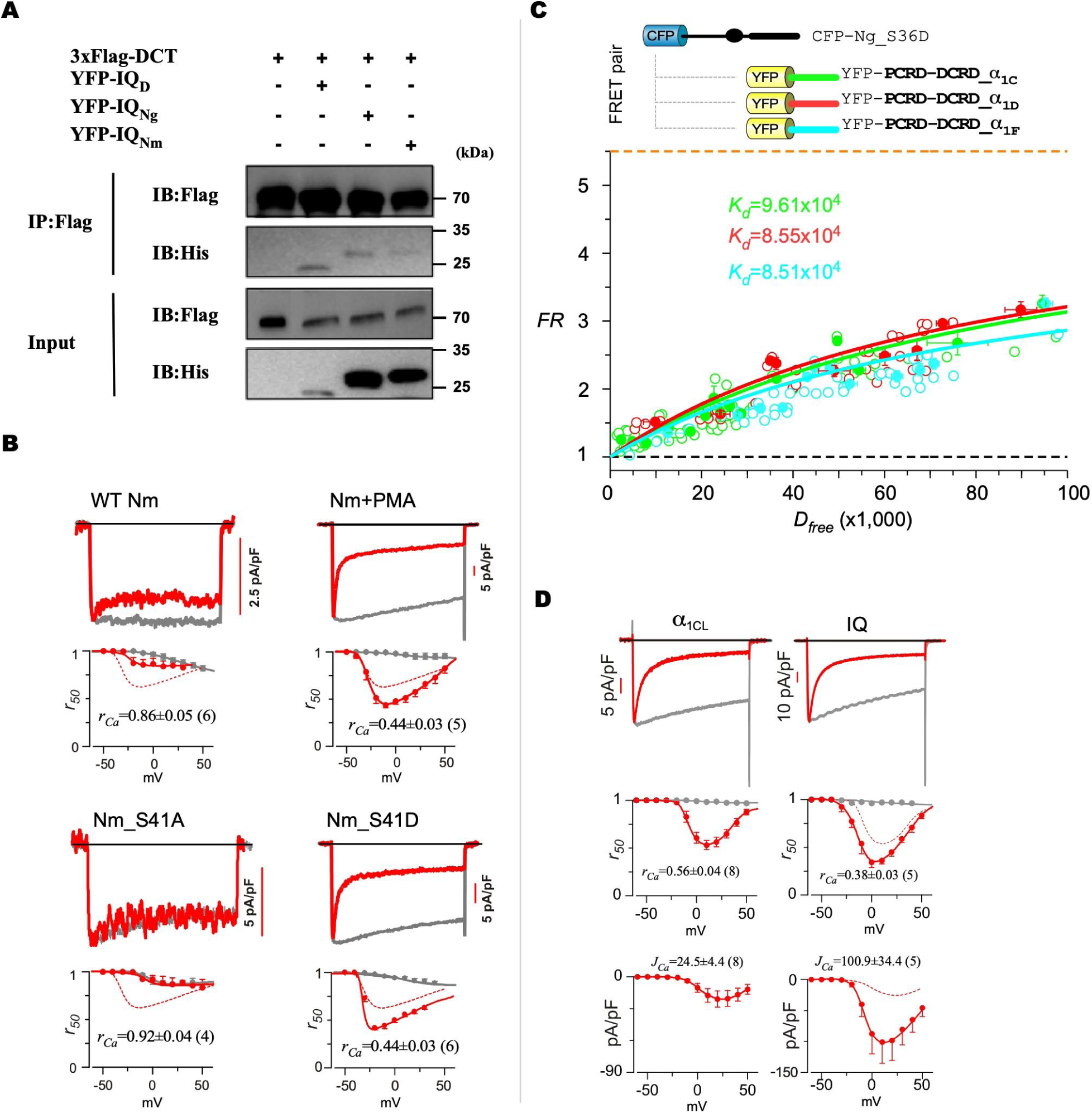
Generalizable interaction and facilitation for more CaM-binding proteins (A) Co-IP results for the potential interactions between DCTD (tagged with Flag) and YFP-IQD, YFP-IQNg, or YFP-IQNm. Cell lysates were immunoprecipitated with anti-Flag magnetic beads, followed by immunoblot analysis. Input lanes represent 10% of the total protein lysate. (B) Electrophysiology for CaV1.3 channels regulated by Nm control, Nm treated with PMA, Nm_S41A, or Nm_S41D. Representative traces and inactivation (*rCa*) profiles are shown. (C) Interactions between CaV1 channel subtypes (PCRD-DCRD_α1C, PCRD-DCRD_α1D, or PCRD-DCRD_α1F) and Ng_S36D were unveiled by FRET 2-hybrid binding assays, indexed by respective *Kd* values. (D) Electrophysiology for CaV1.2 channels regulated by IQ. Representative traces, inactivation (*rCa*) and activation (*JCa*) profiles are shown.

### DCT-bound pNg transfers its unloading CaM to the channel within a transient complex

This new finding of phosphorylation-dependent binding and modulation of CaV1 channels holds significant potential for understanding Ng-related roles in the cell. First of all, our data provide new insights into the classic role of Ng in regulating CaM, the latter of which is well known for its calmodulation role in CaV1 functions. In this context, what would be the benefits for Ng to directly interact with CaV1? We first gained our insights from computational modeling of the structures and interactions of proteins, using AlphaFold3 ^27^ and other tools (see **Materials and Methods**). CaV1.3 adopts two distinct conformational states as manifested by its carboxyl terminus (CT) containing both PCT and DCT. First, the auto-inhibited state is characterized by lower amplitude and weaker CDI overall permitting less Ca^2+^ influx. The simulation results are consistent with the expected conformation that DCT is associated with PCT around the α-helical IQD motif, supported by multiple lines of experimental evidence ^19,21,28^. To achieve the unlock state of the channel, apoCaM is supplied to bind the channel around the IQD domain (**Figure 6A**, top). The CaM C-lobe is primarily engaged with IQD, while its N-lobe interacts with the preIQ region, switching the channel out of its autoinhibition where PCT and DCT is not bound to each other anymore. Notably, AlphaFold3 was unable to produce the structure of an extended CT without adding apoCaM, suggesting that the disinhibited channel or CT is in an unstable or transient state. To further confirm these viewpoints, Co-IP and FRET experiments were conducted showing that apoCaM is able to bind PCT of CaV1.3, at its IQ and preIQ presumed to be targeted by C_CaM and N_CaM, respectively (**Figure 6S1**). Meanwhile, following the experimental settings where a glycine linker was used between Ng and CaM ^29^, the AlphaFold3 modeling generated a dumbbell-shaped complex. The structure indicated the major interface lies between IQNg (the IQ domain of unphosphorylated Ng) and C_CaM (the C-lobe of apoCaM). The binding affinity would be substantially reduced by PKC-mediated phosphorylation at S36 of IQNg. Interestingly, phosphorylation induced the formation of a *de novo* α-helix within the N-terminus of pNg (near the 17^th^ residue), facilitating its connection to N_CaM (the N-lobe). Such structural rearrangement has been reported in CaMKII where the unstructured region in the upstream of the CaM-binding domain is transformed into α-helix upon T286 phosphorylation ^30^. Similar to the above case of PCT/DCT unlocking, with the aid of the new interaction we discovered between DCT and pNg (**Figure 4**), another transient state was unveiled that CaM is bound onto the N-terminus of pNg (**Figure 6A**, bottom). Confirmed by both Co-IP and FRET experiments, Ng in its basal state (Ng_S36A) strongly binds apoCaM; and more specifically, by binding the C-lobe (C_CaM) with minor contributions from the N-lobe (N_CaM), presumed to be mainly at the IQ and preIQ regions of Ng, respectively (**Figure 6S2**). This is in part supported by the fact that the binding affinities of Ng_S36D to CaM or C_CaM were much weaker than Ng_S36A.

**Figure 6.**
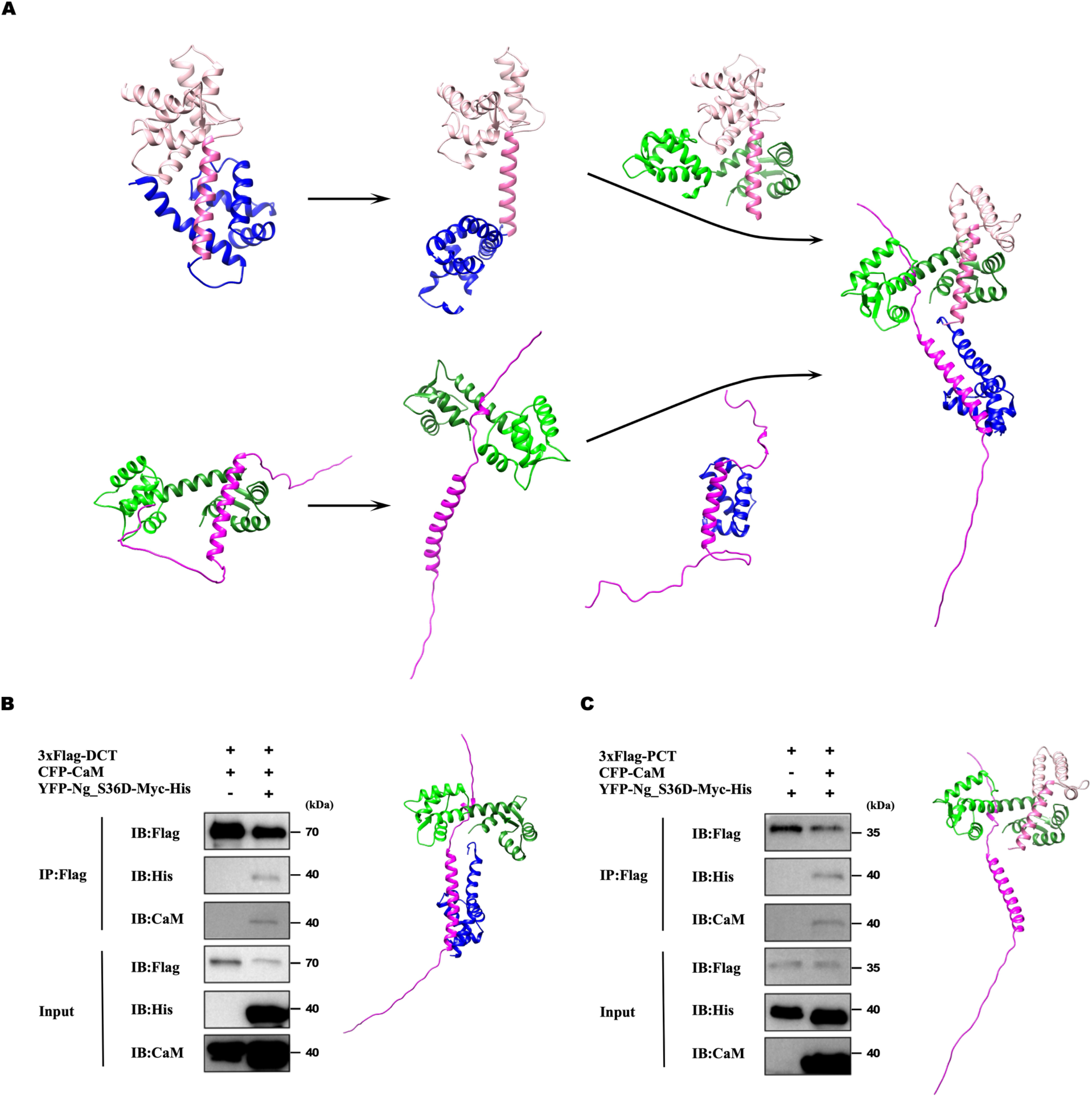
A transient complex of pNg-CaM with CaV1 enables specific CaM transfer (A) Structural models of the CaV1.3 CT, Ng and CaM complex. The CT structure is switched from an autoinhibition state to a disinhibition state (top row), where the DCT (blue) domain is either bound with the PCT region containing the preIQ (colored pale pink) and IQ (pink) domains or unbinding from the preIQ-IQ thereby exposing its PCT and DCT domains (top, middle). apoCaM (green) is known for its binding to preIQ-IQ to promote channel opening, which could compete with autoinhibitory DCT. Structural models of the interactions between Ng (magenta) and CaM (two lobes: C_CaM, dark green; N_CaM, green), where C_CaM is bound at the IQNg domain (bottom row). Upon phosphorylation, C_CaM departs from IQNg, while a transient complex is formed between the newly emerged helical structure at the N-termins of pNg and N_CaM. This transient state was unveiled with the aid of the interaction between DCT and IQNg discovered in this study. Altogether, the whole complex is formed by CT, pNg and CaM (right, middle). pNg binds onto DCT, which unlocks CT autoinhibition while maintaining the connection with N_CaM, thereby allowing C_CaM to be specifically and effectively delivered to the IQD domain of the channel within the signaling complex. (B) Phosphorylated Ng concurrently links CaV1 DCT and CaM, suggested by the Co-IP results on the interactions among CaM, Ng and CaV1.3 DCT. HEK293 cells were transfected with DCT tagged with Flag and CFP-CaM, either alone or in the presence of YFP-Ng_S36D-Myc-His. Cells were subjected to Co-IP 48 hours later using anti-Flag-magnetic-beads, followed by western blot with anti-His antibody, anti-CaM antibody or anti-Flag antibody. Input lanes represent 10% of total protein lysate. Three independent experiments were performed, yielding similar results. The structure partially taken from the whole complex (**A**) illustrates the linkage of pNg to both DCT (DCRD) and CaM (C_CaM). (C) The concurrent linkage of CaM to pNg and CaV1 is shown by Co-IP results on the interactions among CaM, Ng and CaV1.3 PCT. HEK293 cells were transfected with preIQ-IQ- PCRD tagged with Flag and YFP-Ng_S36D-Myc-His, either alone or in the presence of CFP- CaM. Cells were subjected to Co-IP 48 hours later using anti-Flag magnetic beads, followed by western blot with anti-His antibody, anti-CaM antibody or anti-Flag antibody. Input lanes represent 10% of total protein lysate. Three independent experiments were performed and similar results were obtained. The structure partially taken from the whole complex (**A**) illustrates the linkage of CaM to both PCT (preIQ-IQ) and pNg.

Taken together, the interactions between CaM and CaV1 closely resemble those between CaM and Ng, especially for apoCaM.

A closer look was given to the complex of pNg and DCT, the focus of this study. F37 has been identified as a critical residue at the binding interface of IQNg/DCRD, which is also confirmed by the structural prediction here. A glycine linker was introduced to connect PCRD and DCRD (PCRD-Gn-DCRD) before loading into AlphaFold3, yielding an initial structural model that exhibited weak intrinsic interactions between PCRD and DCRD. Subsequent AlphaFold3 co-modeling of pNg with the PCRD-Gn-DCRD construct revealed that pNg stabilizes the PCRD-DCRD interaction by bridging the two domains. Notably, F37 of pNg is confirmed to interact with DCRD as well as PCRD in the complex. These modeling data serve as valuable information in line with the experimental results shown earlier.

Building upon the preceding modeling results, we propose the potential mechanism of the targeted CaM delivery by Ng. A signaling complex was achieved by AlphaFold3 and GRAMM docking, including all the four key components: PCT, CaM, Ng, and DCT (**Figure 6A**, right). pNg plays critical roles with two sets of interactions with both DCT and N_CaM; and CaM is also bound with two partners with both lobes: N_CaM/pNg and C_CaM/PCT. In this scheme, while unloading CaM from IQNg, pNg binds onto DCT of the channel. In this transient state, pNg still connects N_CaM, allowing the preferential binding of the C_CaM on IQD of the channel due to the ultrahigh equivalent concentration or affinity within the same molecule or complex. Effectively, CaM is being transferred from Ng to the CaV1 channel.

Co-IP results provided further support that CaV1.3 DCT, Ng_S36D, and CaM could form a ternary complex. DCT itself could not bind to CaM; however, in the presence of Ng_S36D, both Ng_S36D and CaM bands were detectable using Co-IP, consistent with the predicted structure (**Figure 6B**, left). Similarly, PCT itself did not directly bind Ng_S36D. However, in the presence of CaM, Co-IP results suggested that CaM could jointly link PCT with Ng_S36D, also well matching the modeling results (**Figure 6B**, right).

These results altogether demonstrate that pNg here not only unlocks the channel via pNg/DCT binding, but also delivers CaM to the channel via pNg/N_CaM binding. Such dual roles of pNg would serve as a new type of “active delivery” of CaM (and potentially other key signaling molecules), to a broad scope of autoinhibited functional proteins (e.g., channels and enzymes). The high similarities between CaM and DCT, and also PCT and Ng, two pairs of binding partners stand out, i.e., CaML modules (CaM and CaM-Like DCT) and CaMBD (CaM-binding domain) containing modules (Ng and PCT). If these CaML/CaMBD interactions are interchangeable, the logic of our discovery is even more straight-forward: all the other interactions have been previously proven except the unexplored pair of DCT and Ng. This would imply an extensive list of binding pairs awaiting future investigations, considering the large number of CaM-related proteins that have been identified ^31,32^.

### pNg promotes CaV1 signaling in neurons but not for disease-related Ng mutants

Besides the above mechanistic insights, what would be the physiological and pathological implications of the signaling complex of pNg, CaM and CaV1 unveiled in this study? The reduced levels of Ng expression within pyramidal neurons are closely linked to aging, diminished thyroid function and cognitive deficits ^13,33^. The running hypothesis before this study pointed to that Ng would act as CaM buffers to modulate the activation of post-synaptic Ca^2+^/CaM-dependent signaling by influencing the spatial distribution and the "functionality" of CaM ^9,34^. On one hand, non-phosphorylated Ng is able to reduce [CaM] (the concentration of free CaM), thereby attenuating CaV1 activities, and the downstream transcriptional signaling to the nucleus. On the other hand, for pNg in the context of this work, CaV1 signaling should be upregulated through the dynamic interactions among pNg, CaM and CaV1.

To elucidate the role of pNg across developmental stages (in DIV, days *in vitro*), we conducted DIV- dependent morphological and functional analyses of cultured cortical neurons expressing Ng_S36D compared to controls, both infected with GCaMP6m-XC to enable long-term imaging of Ca^2+^ oscillations (**Figure 7A**) and neuritogenesis (**Figure 7B**). Neurite tracing revealed substantial enhancement in the total length of neurites, while concurrent Ca^2+^ imaging demonstrated the changes in the amplitudes and frequency of the autonomous Ca^2+^ oscillations (**Figure 7S1**). As shown in previous studies, CaV1 activities and neuritogenesis are coupled with each other, which could be quantified by a phase-plane-like trajectory between AUC (Area Under the Curve representing the Ca^2+^ influx) and NGR (Neurite Growth Rate) ^35,36^. With Ng_S36D overexpression, AUC was overall enhanced in comparison to controls, as evidenced by an upward shift in the bell-shaped curve; similarly, NGR was elevated, following a parallel enhancement pattern (**Figure 7C**). Phase-plane analysis of AUC-NGR revealed distinct trajectory patterns between control and Ng_S36D-expressing neurons. In Ng_S36D neurons (pink curve), the trajectory was shifted rightward and upward relative to controls (black curve), indicating a global enhancement of both Ca²⁺ influx and neuritogenesis. The dynamic range was expanded under Ng_S36D expression, with higher peak AUC and NGR values and a prolonged growth phase. These observations support the notions that pNg (but not CaM-occupied Ng without phosphorylation) promotes CaV1 gating (shown earlier) and its downstream signaling (shown here). As stated, the role of pNg is to deliver CaM to the channel, which effectively elevates the (local) concentration of CaM. We performed the CaM-overexpression experiments and the results are consistent with pNg (**Figure 7S2**). As expected, both autonomous oscillations and neurite outgrowth were enhanced by CaM up to DIV14, supposedly in a concentration-dependent manner ^19^. In physiological conditions, it is unrealistic to have so high supply of CaM as in the overexpression experiment. More physiological scenario could be related to the mechanisms and roles of pNg unveiled in this work, which serves as a phosphorylation/activity-dependent scheme for CaV1 and its signaling to achieve the goal of targeted CaM-tuning or calmodulation.

**Figure 7.**
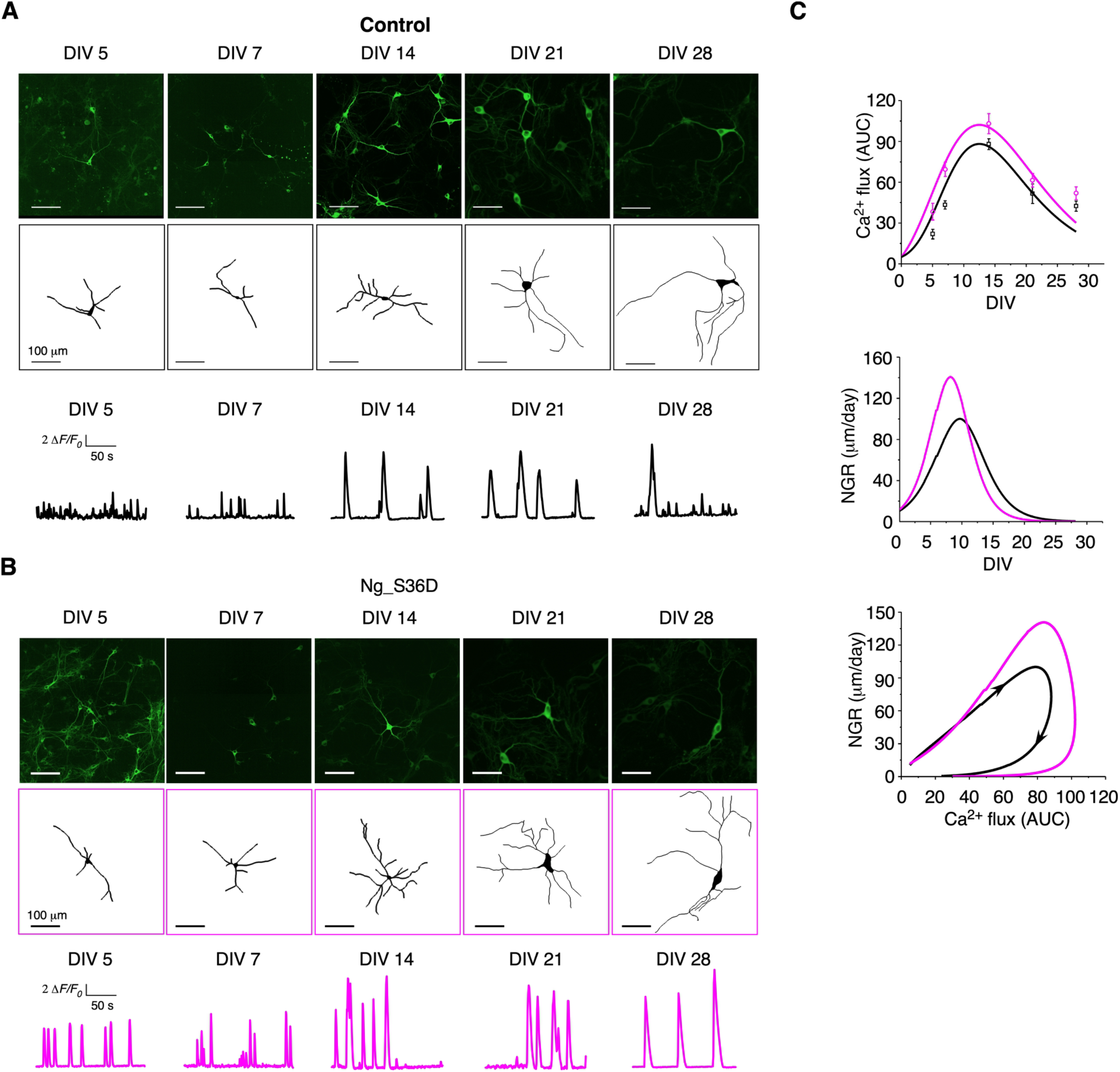
pNg in cooperation with CaM facilitates CaV1 unlocking and signaling (**A**-**C**) Effect of Ng_S36D on the Ca^2+^ influx-neuritogenesis coupling in cortical neurons. Neurite tracing (**A**) and fluorescence Ca^2+^ imaging (**B**) from the control neurons versus the neurons expressing Ng_S36D all infected with GCaMP6m-XC viruses, summarized in (**C**). Time-dependent profiles of the peak amplitude (*ΔF/F0*). The temporal profiles of the average frequency of Ca^2+^ oscillations. Time-dependent profiles of Ca^2+^ influx, described by the correlation between AUC (*ΔF/F0* per min) of Ca^2+^ oscillations and the development stage (DIV). Time-dependent profiles of neurite outgrowth, described by the correlation between the total length per single neuron (μm) and the development stage (DIV). Temporal profiles of neurite growth rate (NGR; μm per day), which is derived from the total length-DIV curve. Relationships between AUC and NGR, the arrows indicate the direction of time. All data points are presented as Mean ± SEM.

Ng is also implicated in Alzheimer’s Disease (AD) that the patients exhibit elevated concentrations of neurogranin in cerebrospinal fluid (CSF) ^37^, a valuable biomarker for early AD diagnosis ^38^. Ng polymorphism and mutants are potentially pathogenic ^39,40^. However, the mechanisms underlying Ng correlations with neural degeneration have remained elusive. We here explored the potential clues in the context of the emerging scheme from this work. Disease-related truncation sites of Ng are primarily located at amino acids 33, 40, and 42 within Ng, situated within its IQ domain ^40^ (**Figure 8A**). By Co-IP, the results demonstrated that the two major mutants of 1-40 and 1-42 lost the key feature of pNg, as compared to the strong affinity between Ng_S36D and DCTD (**Figure 8B**). Also, the Ng mutants failed to regulate CaV1.3 functions as compared to the effects of Ng_S36D on channel activation and inactivation (**Figure 8C**). In cortical neurons, the faciliatory effects of pNg was diminished by Ng mutations, as evidenced by significantly less Ca^2+^ influx and neurite outgrowth in neurons transfected with Ng_S36D_1-40 and Ng_S36D_1-42 (**Figure 8D-F**). Comparable findings were obtained from hippocampal neurons, where oscillations play a vital role in learning and memory (**Figure 8S1**).

**Figure 8.**
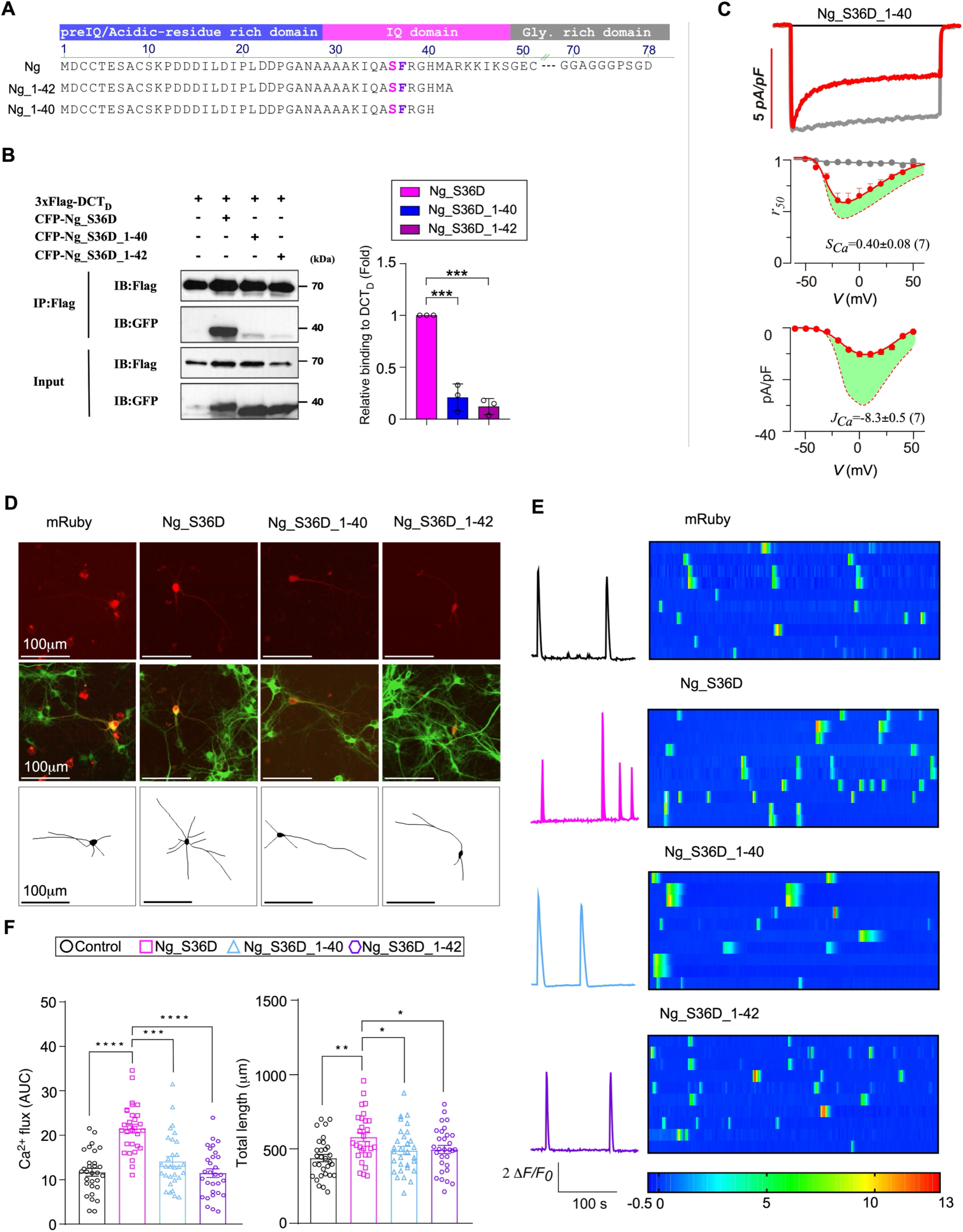
Disease-related mutations diminished the facilitation effects of pNg on cortical neurons (A) The predominant truncation mutants of Ng_S36D: Ng_S36D_1-40 and Ng_S36D_1-42. Green shades indicate the differences from the control (Ng_S36D). (B) Co-IP to examine the interactions between DCTD (tagged with Flag) and CFP-Ng_S36D, CFP-Ng_S36D_1-40 or CFP-Ng_S36D_1-42. Cell lysates were immunoprecipitated with anti- Flag magnetic beads, followed by immunoblot analysis. Input lanes represent 10% of the total protein lysate. (C) Electrophysiology for CaV1.3 channels overexpressed with Ng_S36D_1-40. Representative traces, inactivation (*rCa*) and activation (*JCa*) profiles are shown. (D) Neurite tracing for the control neurons versus the neurons expressing Ng_S36D, Ng_S36D_1-40 and Ng_S36D_1-42. (E) Calcium oscillations are depicted by the trace of one representative neuron (left) and the heat map from multiple neurons (right). (F) Two key indices of AUC and total neurite length were calculated and compared for the Ng_S36D control versus the truncation mutants. All statistical data are presented as Mean ± SEM. One-way ANOVA followed by Dunnett for post hoc test for (F): *, *p* < 0.05; **, *p* < 0.01; ***, *p* < 0.001; n.s., *p* > 0.05.

## Discussion

In this work, we unveil an unprecedented inter-relationship between Ng, CaM and CaV1 (**Figure 9**). It has been well documented that both Ng and CaV1 bind apoCaM with high affinities at their IQ domains. Unexpectedly, phosphorylated Ng (pNg) directly binds CaV1 DCT. Meanwhile, the CaM being unloaded from pNg would be favored (over ambient CaM) to deliver onto CaV1 PCT. Taken together, Ng and CaM cooperate as an activity/phosphorylation-dependent complex to unlock CaV1 autoinhibition and facilitate CaV1 signaling.

**Figure 9.**
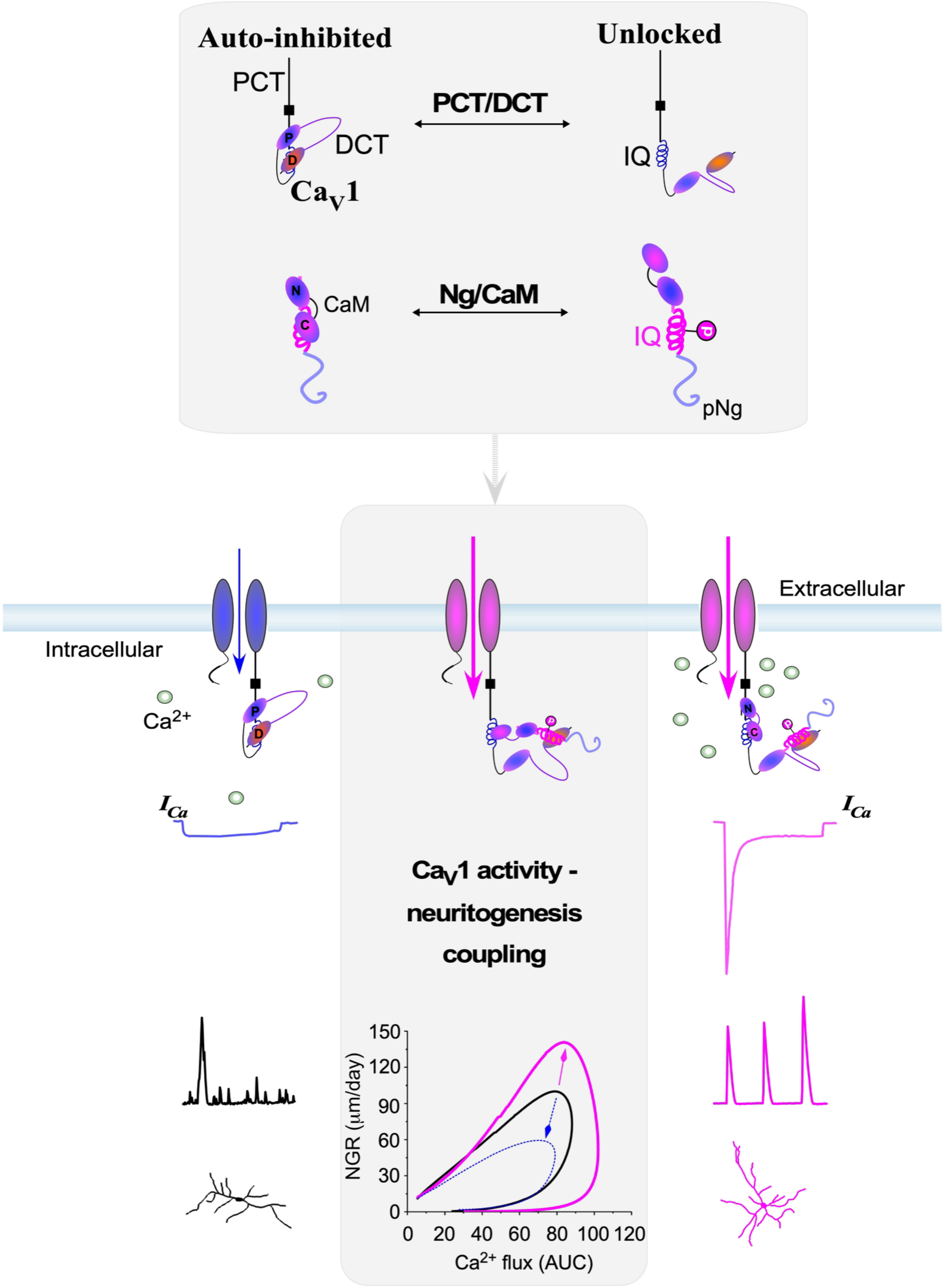
Ng unlocks CaV1 autoinhibition and promotes CaV1 signaling PCT/DCT and Ng/CaM share high similarities in dynamic interactions (top, gray area). For autoinhibited CaV1 CT, PCT (containing IQ) and DCT (containing PCRD and DCRD domains, denoted as P and D) are tightly bound, similar to CaM-buffering Ng. When phosphorylated Ng is unloading CaM, pNg essentially forms a (transient) complex with CaM, similar to the unlocked CT, where CaM-binding domain preIQ-IQ is unbound with CaM-like domain DCT. The dynamic interactions between PCT/DCT and Ng/CaM promote CaV1 gating and signaling (lower illustration), switching CaV1 from the autoinhibited state (smaller CaV1 current, weaker Ca^2+^ oscillation and less growth) to the unlock state (larger CaV1 current, stronger Ca^2+^oscillation and more pronounced neuritogenesis). The central mechanism (highlighted in grey) is the complex formed by the direct interactions between Ng (IQNg) and CaV1 (DCT), where the pNg releases its CaM to the CaV1 channel, thereby effectively and specifically promoting (pink) the CaV1 activity-neuritogenesis coupling in direct contrast to the disease-related Ng mutants (dotted blue).

### Intermolecular interactions between CaM (or like) and CaM-binding (or like) domains

Ng, a member of calpacitin proteins, undergoes regulation through phosphorylation by PKC ^10^ and dephosphorylation by phosphatase ^41^. In its non-phosphorylated state, Ng demonstrates a strong binding affinity with CaM, which is significantly reduced in response to particular conditions such as high Ca^2+^, phosphorylation, or oxidation ^25,42,43^. Beyond these, no endogenous protein has been reported to directly interact with Ng. Our research has revealed CaV1 as a novel binding target of Ng that phosphorylated Ng or CaM-free Ng directly binds to CaV1.3 DCT. The targets of Ng can expand onto other CaV1/DCT variants (**Figure 5**), and potentially an even broader scope of CaM-like motifs or proteins which await future investigations ^22^. Regarding CaV1, the PCT domain binds CaM at its preIQ-IQ motif, which is competed by its DCT domain ^21^, underlying CaV1 autoinhibition or CMI (C-terminus mediated inhibition) ^6,44–46^. C-terminus is expected to mediate diverse physiological modulations. DCT is reportedly subject to phosphorylation thus presumably relieving the autoinhibition, as suggested by CaV1.2 DCT with PKC ^47^ or PKA ^48^ phosphorylation and by CaV1.4 DCT with PKA phosphorylation ^49^. The binding partners of CaV1 DCT thus far are mostly scaffold proteins such as Densin ^50^, Shank ^51^ anchoring at its very distal PDZ-binding domain, or AKAP binding with a leucine zipper motif of distal DCT ^52^. However, no report has associated any direct binding partner of DCT with the mechanism of disinhibition or activation/facilitation. We here report the first of its kind: Ng, once phosphorylated, forms the direct interaction with DCT via its IQNg domain, immediately expandable onto other calpacitin members (**Figure 5**), and presumably onto even more proteins containing CaM-binding domains ^53^, yet to be identified. In brief, this work opens up new avenues to discover dynamic interactions between CaM (or CaM-like) and CaM-binding (or CaM binding domain-like) motifs, among the vast number of Ca^2+^ signaling proteins.

### New mechanism of CaV1 facilitation

Activity or Ca^2+^ dependent facilitation of CaV1 is in close relevance to critical pathophysiological processes in the heart and brain. Currently, there exist three main mechanisms to enhance or facilitate CaV1 activation: small-molecule agonists, modulatory proteins, and channel phosphorylation. First, Bay K8644 as a widely used CaV1 agonist upregulates the open probability of the channel, by binding to the same site as DHP antagonists, but causing opposite effects ^54,55^. Second, activation of the channel is achieved through phosphorylation, by the kinases of CaMKII, PKC, or PKA ^47,48,56–59^. Third, activation is mediated by specific protein-protein interactions. Rad could reduce its affinity for auxiliary β-subunits, thereby alleviating the constitutive inhibition of CaV1.2 in response to β-adrenergic stimulation or PKA activation (instead of PKA phosphorylation of the channel) ^60^. Most of other studies focus on the interactions with the IQ and its vicinity, by CaM, CaBP, α-actinin, CaMKII ^5,57,61,62^. This work has unveiled a distinct mechanism of CaV1 facilitation highlighted by the direct interactions between CaV1 DCT and phosphorylated Ng. To our knowledge, this is the first report regarding the Ca^2+^/activity/phosphorylation-dependent binding of DCT as the core mechanism driving CaV1 into facilitation or out of autoinhibition. More CaV1 agonists are expected to emerge after this work, potentially from the superfamilies of CaM-binding or like proteins, including but not limited to calpacitin (**Figure 5**). Upon dephosphorylation, Ng, due to its strong affinity with apoCaM, reverts to an inactive state or DCT-unbinding state, thereby CaV1 subsequently returns back to the baseline activities from facilitation. How does this correlate with the potential roles of Ng and CaV1 in activity-dependent plasticity, e.g., long-term facilitation (LTP) ^9,12,34^? The questions along this line, e.g., whether and which DCT-binding proteins would be closely involved in CaV1- associated plasticity, are worth exploring in future research.

### Active and specific CaM delivery to the targeted proteins beyond CaM buffering

Ca^2+^/CaM binds to numerous targets, including protein kinases, phosphatases, membrane channels and receptors, and nitric oxide synthases, leading to diverse signaling pathways ^53,63,64^. In the absence of Ca^2+^, apoCaM also binds to a good number of proteins like myosin-binding proteins, cytoskeletal proteins, enzymes, and ion channels and membrane receptors ^65^. Due to its extensive interactions and limited availability, free (unbound) CaM can be viewed as a common intracellular currency—much like Ca²⁺—that integrates and distributes signaling across diverse cellular pathways ^66^. Based on the effects of Ca^2+^ and phosphorylation on CaM binding, Ng is presumed to serve as a transport vehicle for CaM ^9^. Within dendritic spines, Ng localizes CaM in proximity to the plasma membrane, promoting CaMKII activation at the same site, thereby lowering the threshold for LTP. However, currently no molecular detail has been available as experimental evidence regarding how Ng particularly mediates these processes beyond its elusive role as a CaM buffer, except for some computational attempts and hypothetical insights ^16,17^. In this study, we propose a dynamic binding mechanism featured with active delivery and high specificity (**Figure 6**). Collective evidence suggests that when phosphorylated Ng binds DCT after the C-lobe of CaM leaves its IQ domain, the N-lobe is still able to link with Ng before full detachment.

This transient linkage of CaM provides the preference of this particular CaM (over ambient CaM) to bind/transfer onto the preIQ-IQ domain, the established pre-association site on the channel for apoCaM ^5^. In other words, Ng binds and unloads its CaM to the same CaV1 channel, altogether fulfilling the task of switching CaV1 out of autoinhibition. Meanwhile, it is worth noting that further experimental evidence will be needed to confirm the transient complex formed by CaM, phosphorylated Ng, and CaV1 CT, by choosing appropriate techniques for transient protein-protein interactions ^67^. Beyond Ng and CaV1.3, we postulate that such CaM delivery paradigm may apply to other CaM-binding proteins. For instance, Ng could directly unload CaM to CaMKII ^68–70^, which in turn might act similarly to transfer its CaM onto the downstream targets of CaMKII ^71,72^.

### Pathophysiological implications of the signaling complex of Ng, CaM and CaV1

The reduced levels of Ng expression within pyramidal neurons are closely linked to aging and cognitive deficits ^13,33^. It has been believed that Ng modulates the activation of post-synaptic Ca^2+^/CaM-dependent signaling by influencing the distribution and "function" of CaM, thereby exerting a temporal and spatial impact on synaptic transmission ^9,34^. In the context of this work, the bidirectional roles of Ng in neurons are closely related to phosphorylation-dependent Ng/CaM regulations of CaV1, contributing to the synaptic plasticity. Non-phosphorylated Ng diminishes CaV1 activity, attenuating the excitation-transcription coupling. More broadly, calpacitin and other CaM-binding proteins could modulate the downstream targets beyond CaV1, in similar fashions. Ng is also implicated in various pathological conditions, including traumatic brain injury ^73^, cognitive dysfunction ^13^, intellectual impairment ^74^, schizophrenia ^75^, and Alzheimer’s Disease (AD) ^37^. Emerging as a valuable biomarker for early AD diagnosis ^38^, AD patients exhibit elevated concentrations of neurogranin in cerebrospinal fluid (CSF) ^37^. Moreover, our study gained mechanistic insights into the pathogenic Ng polymorphism and its mutants ^39,40^, serving as a foundation for future investigations, such as animal models and patient-based studies. Meanwhile, human mutations on CaV1 have been found their implications in various diseases, known as CaV1 channelopathy, including neurogenerative diseases ^76^, Timothy syndrome ^77^, autism and intellectual disabilities ^78^, sinoatrial node dysfunction and deafness (SANDD) syndrome ^79^, and adrenal hypertension ^80^. For both Ng and CaV1, the known mutations/truncations include those near their binding interfaces. For instance, Ng truncation sites are primarily located at amino acids 33, 40, and 42 within Ng, situated within its IQ domain ^40^. And regarding CaV1, the C-terminus is subject to mutations, truncations or splicing ^81,82^. In light of this study, DCT mutations could disrupt its binding with Ng and like proteins, consequently causing dysregulations of CaV1 gating and signaling, including gene transcription and synaptic plasticity. The findings and related tools originated from this and follow-up studies would facilitate our understanding and therapeutic interventions for neurodegeneration and other brain diseases.

## Materials and Methods

### Molecular biology

The plasmids of channels and peptides were constructed from α1C (human CaV1.2, AF465484.1), α1D (human CaV1.3 α1DL, NM_000720, rat CaV1.3 α1DL, NM_001389225.2), and α1F (human CaV1.4, NM_005183.4). α1DL-3xFlag was produced previously ^6,83^. CFP/YFP-tagged constructs following the I-II loop, II-III loop, III-IV loop, IQ, PCRD-DCRD, PCT, NT, or CT in pcDNA3 were made using a similar process as described previously ^83^. In brief, the tags of fluorescent proteins (FPs, including CFP or YFP) were cloned into the pcDNA3 vector by KpnI and NotI, then desired segments were subcloned to the C-terminus of FP via the unique NotI and XbaI sites. 3xFlag-DCTC, 3xFlag-DCTD, 3xFlag-DCTF, 3xFlag- preIQ-IQ-PCRD, and 3xFlag-PCT-DCT were generated in the same way: the tag of 3xFlag (DYKDHDGDYKDHDIDYKDDDDK) was cloned into pcDNA3 vector by KpnI and NotI, then aim fragments were subcloned via the unique NotI and XbaI sites. Point mutations on YFP-IQD including IQ_I/A, IQ_Q/A, IQ_F/A, IQ_R/A, IQ_FR/AA, IQ_K/A, and IQ_RK/AA were made using Mut Express MultiS Fast Mutagenesis Kit V2 (Vazyme).

For neurogranin (Ng) and neuromodulin (Nm) related constructs, Nm (NM_017195.3) and Ng (NM_024140.3) were cloned from a rat brain cDNA library and inserted into the vector pIRES2-EGFP via BglII-SacII sites or YFP-insert/pcDNA3 via NotI and SpeI, respectively. YFP-Ng_S36A, YFP-Ng_S36D, Nm_S41A (IRES-GFP), and Nm_S41D (IRES-GFP) were made using QuikChange Site-Directed Mutagenesis Kit (Agilent Technologies/Vayzme) or overlap PCR. CFP-tagged constructs (CFP-Ng_S36A, CFP-Ng_S36D, CFP-Ng_S36D_F37A, CFP-Ng_S36D_FR/AA) were made in the same way mentioned above. For subsequent Co-IP related experiments, YFP-Ng, YFP-Ng_S36A, and YFP-Ng_S36D were cloned into the pcDNA3.1-Myc-His A vector by NotI and EcoRI. Three domains of Ng: preIQNg, IQNg, preIQ-IQNg, and GRDNg were cloned into the 3xHA-pEGFP-N1 vector by HindIII and KpnI. IQNg_F13A was synthetized through Tsingke company.

The tag of CFP was cloned into the pcDNA3 vector by KpnI and NotI, then desired segments (CaM, C_CaM, N_CaM) were subcloned to the C-terminus of CFP via the unique NotI and XbaI sites. CFP-CaM- 3xFlag, CFP-C_CaM-3xFlag, and CFP-N_CaM-3xFlag were generated in the same way: the tag of 3xFlag (DYKDHDGDYKDHDIDYKDDDDK) was fused to the C-terminus of then desired segments. YFP-IQNg- G12-C_CaM was synthetized through Tsingke company. Flag-C_CaM, Flag-N_CaM, and Flag-IQNg-G12- C_CaM were cloned in the same manner as described above.

### Transfection of cDNA constructs

In this study the HEK293 cell line (ATCC) was free of mycoplasma contamination, checked by PCR with primers 5′-GGCGAATGGGTGAGTAACACG-3′ and 5′-CGGATAACGCTTGCGACCTATG -3′ ^19,83^. For electrophysiology, HEK293 cells were cultured in 60 mm plates, and the constructs, including those of channels, were transiently transfected using the calcium phosphate protocol. We applied 4-5 μg of cDNA encoding the α1 subunit of the channel, along with 4 μg of rat brain β2a (NM_053851.2) and 4 μg of rat brain α28 (NM012919.3) subunits. Additional cDNA constructs were added during transfection when necessary. To enhance expression, cDNA for simian virus 40 T antigen (1 μg) was co-transfected. For all the construct transfections, 2 μg of plasmids were added together with the channel constructs. After 6-8 hours, the dishes were washed 2-3 times with pre-warmed PBS, and complete medium was added for culture. Electrophysiological experiments were performed at room temperature 2 days after transfection.

For FRET assays and Co-Immunoprecipitation assays, transfections of plasmids into HEK293 cells were carried out by using Lipofectamine 2000 (Invitrogen). Briefly, plasmids and transfection reagent (1.5 μl per μg of DNA) were each diluted with Opti-MEM (Invitrogen), mixed together, and incubated for 20 minutes at room temperature. The mixture was then added to the medium for transfection. After 6-8 hours, the medium was replaced by serum medium. Cells were used at 24 to 48 hours after transfection.

### Whole-cell electrophysiology

Patch-clamp whole-cell recordings of transfected HEK293 cells were performed at room temperature using an Axopatch 200B amplifier (Molecular Devices). Electrodes were pulled with borosilicate glass capillaries by a programmable puller (P-1000, Sutter Instruments) and heat-polished using a microforge (MF-830, Narishige), resulting in 2-5 MΩ resistance. Series resistance was compensated by 70%. Whole-cell currents were generated from a family of step depolarizations (−70 to +50 mV from a holding potential of −70 mV with a step increase of 10 mV), according to our previous protocol ^6^. The internal solutions contained (in mM): CsMeSO3, 135; CsCl2, 5; MgCl2, 1; MgATP, 4; HEPES (pH 7.3), 5; and EGTA, 5 at 290 mOsm, adjusted with glucose. The bath solution contained (in mM): TEA-MeSO3, 140; HEPES, 10 (pH 7.3); CaCl2 or BaCl2, 10; 300 mOsm, adjusted with glucose, similar to the previous protocols ^83^.

### FRET optical imaging

FRET two-hybrid experiments were carried out on an inverted epi-fluorescence microscope (Ti-U, Nikon), with computer-controlled filter wheels (Sutter Instruments) to cooperate with dichronic mirrors for both excitation and emission via each imaging channel. For CFP/YFP-based FRET experiments, the following filters were used: excitation: 438/24 nm (FF01-438/24-25, Semrock) and 480/30 nm (FITC, Nikon); and emission: 483/32 nm (FF01-483/32-25, Semrock) and 535/40 (FITC, Nikon). The dichroic mirrors used were: 458 nm (FF458-Di02-25x36, Semrock) and 505 nm (FITC, Nikon). The actual power of excitation was in the range of 5-15 mW at the site of specimens, as measured by the optical power meter (PM-100, Thorlabs).

Fluorescent images were acquired by a Neo sCMOS camera (Andor Technology) and analyzed with 3^3^- FRET algorithms coded in Matlab (Mathworks) ^19,84^, based on

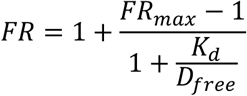

where *FRmax* represents the maximum FRET ratio *FR* pertaining to the receptor (YFP), and *Dfree* denotes the equivalent free donor (CFP) concentration. By fitting the curve of *FR-Dfree* with a set of customized Matlab codes to iteratively estimate *FRmax* and *Dfree*, the values of the effective dissociation equilibrium constant (*Kd*) can be achieved for each binding pair to compare the relative affinity. FRET imaging was performed with HEK293 cells in Tyrode’s buffer containing 2 mM Ca^2+^ or PBS solution.

### Western blot and Co-Immunoprecipitation

HEK293 cells were transfected using Lipofectamine and cultured for 2 days before cell lysates were prepared with lysis buffer RIPA (cat#P0013C, Beyotime) (with protease inhibitor cocktail, cat#P1006, Beyotime) and centrifugation at 14,000×g for 5 min at 4°C. 10% of the supernatant was individually extracted as respective input samples. The supernatants were incubated with 30 μl of anti-Flag Magnetic Beads (cat#B26101, Bimake) overnight at 4°C for Co-Immunoprecipitation. The proteins were separated using 1xSDS-PAGE Protein Sample Loading Buffer (cat#P0286-15ml, Beyotime). The washed samples were boiled at 100°C for 8 min (until complete denaturation of proteins), followed by a 2 min incubation on ice. Then western blotting was performed using the indicated antibodies.

For western blotting, equal amounts of protein were separated using 10% sodium dodecyl sulfate polyacrylamide gel electrophoresis and transferred to a nitrocellulose membrane for 20-40 min using rapid transfer Buffer (NcmBlot Rapid Transfer Buffer, cat#WB4600, NCM Biotech). After that, the membrane was blocked in 5% nonfat dry milk for 2 hours at room temperature. Then the nitrocellulose membrane was incubated with primary antibodies (polyclonal anti-Flag antibody, cat#20543-1-AP, Proteintech, Source: Rabbit, Dilution: 1:5000; monoclonal anti-GFP antibody, cat#66002-1-Ig, Proteintech, Source: Mouse, Dilution: 1:10000; His-Tag mAb, cat#P01L071, Gene-Protein Link Biotech, Source: Rabbit, Dilution: 1:1500; Monoclonal anti-Calmodulin antibody, cat#EP799Y, Abcam, Source: Rabbit, Dilution: 1:10000) overnight at 4°C. Next, the membrane was washed three times with TBST and incubated with the appropriate secondary antibody (Goat anti-Mouse, cat#SA00001-1, Proteintech, Dilution: 1:10000; Goat anti-Rabbit, cat#SA00001-2, Proteintech, Dilution: 1:10000) for 2 h at room temperature and then washed three times with TBST again. Finally, the membrane was covered with ECL chemiluminescent reagent (cat#P0018M, Beyotime). The protein levels were detected by an enhanced chemiluminescence system. Three or more replicates were performed for each sample as required.

### Computational Simulations of Protein Structures

We generated predicted structural models for CTD (CaV1.31479-2161), DCRDD (CaV1.32045-2107), IQD (CaV1.31587-1611), CaM, Ng, pNg and IQNg (Ng27-48) proteins using the AlphaFold3 (AF3) platform ^27^. Subsequently, the AF3-Multimer algorithm was employed to model their interaction (CT-mediated Autoinhibition, Complex of CT and apoCaM, pNg and DCRD). Fusion sequences of Ng/pNg and CaM were implemented in AF3 structural modeling to predict the Ng/pNg-CaM complexes. The structures of IQNg/DCRDD and IQD/DCRDD were predicted by the online program of GRAMM ^85–87^ (DCRDD by PyRosetta and I-TASSER). To model the structure of pNg-mediated delivery of CaM to CaV1.3, we extracted the high-scoring structure for CT. PCRD and DCRD were linked with G10 (instead of the long flexible region in between, Figure 2S2) to represent DCT for more efficient modeling. Protein models were evaluated based on interaction scores to find the most plausible binding configuration by GRAMM.

### Dissection and culturing of cortical and hippocampal neurons

The brains were isolated from newborn ICR mice or SD rat (sex of animals not considered as a variable in this study), and cortical neurons or hippocampal neurons were dissected, respectively. The tissues of cortical neurons or hippocampal neurons were incubated in 0.25% trypsin without EGTA for 15 min at 37°C. The tissue digestion suspension was terminated using DMEM supplemented with 10% FBS and 1% antibiotics. The clumps were removed by filtering.

The cell suspension was centrifuged at 1,000 rpm for 5 min, and the cells were resuspended in DMEM containing 10% FBS, then plated onto poly-D-lysine-coated 35 mm confocal dishes. After 4 h, the medium was replaced with Neurobasal medium supplemented with 2% B27 and 1% GlutaMAX-I (growth medium), and maintained in an incubator with a temperature of 37°C and 5% CO2. All animals were obtained from Beijing Vital River Laboratory Animal Technology Co., Ltd. Procedures involving animals were approved by local institutional ethical committees (Beihang University).

### Virus infection on cultured neurons

AAV2/DJ-hSyn-jGCaMP7b-XC virus was used for infection of cultured neurons (Hanbio Biotechnology, China). Other viruses include pLenti-hsyn-mcherry-WPRE, pSLenti-hSyn-mRuby-Ng_S36D-WPRE (OBiO Technology, China). 1 mL of 1 x 10^12^ v.g./mL of the desired adeno-associated virus or 1–2 mL of 1 x 10^8^ TU/ml of the desired lentivirus were added to growth medium at DIV 0 unless otherwise indicated. Neuronal experiments were repeated independently at least twice.

### Transfection of cDNA constructs in neurons

The neuronal culture medium was replaced with pre-warmed Neurobasal medium. 1 μg of cDNA was transiently transfected into DIV 3–5 cultured neurons by Lipofectamine 2000 (Invitrogen) with a typical protocol according to the manual. The Opti-MEM containing plasmids and Lipofectamine 2000 was added to the culture medium for transfection. After 2 h, the culture medium was replaced with Neurobasal medium supplemented with 2% B27 and 1% GlutaMAX-I, and the cells were cultured for at least 2 days before imaging.

### Ca^2+^ imaging with GCaMP-X in cultured neurons

Ca^2+^ imaging of neurons expressing GCaMP-X was performed by Dragonfly High Speed Confocal Microscope (Dragonfly 200, Andor, England) and with Fusion software. Fluorescence intensity (F) was subtracted from its background. F0 is the baseline fluorescence averaged from five data points at rest, and *ΔF* = *F-F0*. *ΔF/F0* serves as the index for Ca^2+^ dynamics. Ca^2+^ waveforms were analyzed by the gadget of Quick Peaks in Origin software with the Three-Standard-Deviations Rule (values >3 SD). AUC (Ca^2+^ influx) was obtained by integrating the area under the curve using Origin software.

Neurite tracings were depicted and measured with Imaris 7.7.2 (Bitplane) from the images under average intensity projection. Only non-overlapping neurons were selected for analysis and images of at least 21 neurons from two independent culture preparations were analyzed.

### Data analysis and statistics

Data were analyzed in Matlab, Origin, and GraphPad software. The standard error of the mean (S.E.M.) and unpaired Student’s *t*-test (two-tailed with criteria of significance: *, *p*<0.05; **, *p*<0.01 and ***, *p*<0.001), unless indicated otherwise, were calculated when applicable.

## Acknowledgements

We thank all X-Lab members for discussions and help. Drs. Min Liu, Yaxiong Yang and Nan Liu conducted pilot experiments for this project. This work is supported by National Natural Science Foundation of China (32371301).

## Author Contributions

XDL conceived the project. Both ZY and JLG made significant and critical contributions. SQ and WZ participated in the experiments and analyses. MJH and YZ conducted simulations and analyses for the complex of Ng/CaM/CaV1. XDL and ZY wrote and finalized the manuscript. All authors contributed to writing and revising the manuscript.

