## Supplementary Figures for "Neurogranin Unlocks the L-type Ca²⁺ Channel and Directs Calmodulin Delivery"

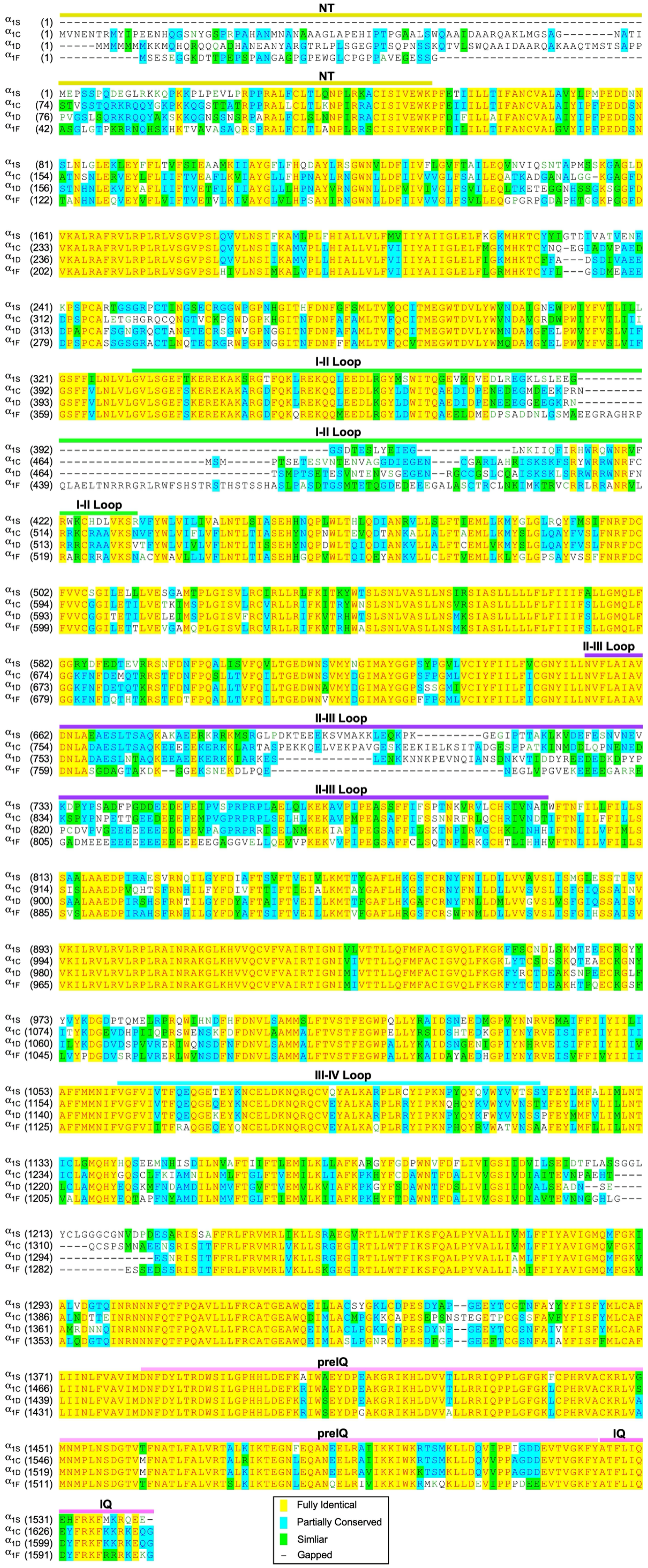

**Supplementary Figure 2S1. Alignment of the channel sequences (up to the IQ domain) across the Ca$_V$1 family**

The channel sequences including the domains of NT, I-II Loop, II-III Loop, III-IV Loop, preIQ and IQ were aligned for Ca$_V$1.1 ($\alpha_{1S}$, NM_001101720.1), Ca$_V$1.2 ($\alpha_{1C}$, AF465484.1), Ca$_V$1.3 ($\alpha_{1D}$, EU363339.1) and Ca$_V$1.4 ($\alpha_{1F}$, NP005174), with GenBank accession numbers in parentheses. Key domains of NT (yellow), I-II Loop (green), II-III Loop (purple), III-IV Loop (cyan), preIQ (pink) and IQ domain (hot pink) are indicated with solid lines in different colors. Sequence homology from high to low is categorized as yellow (fully identical), cyan (partially conserved), or green (similar) and – (gapped or different).

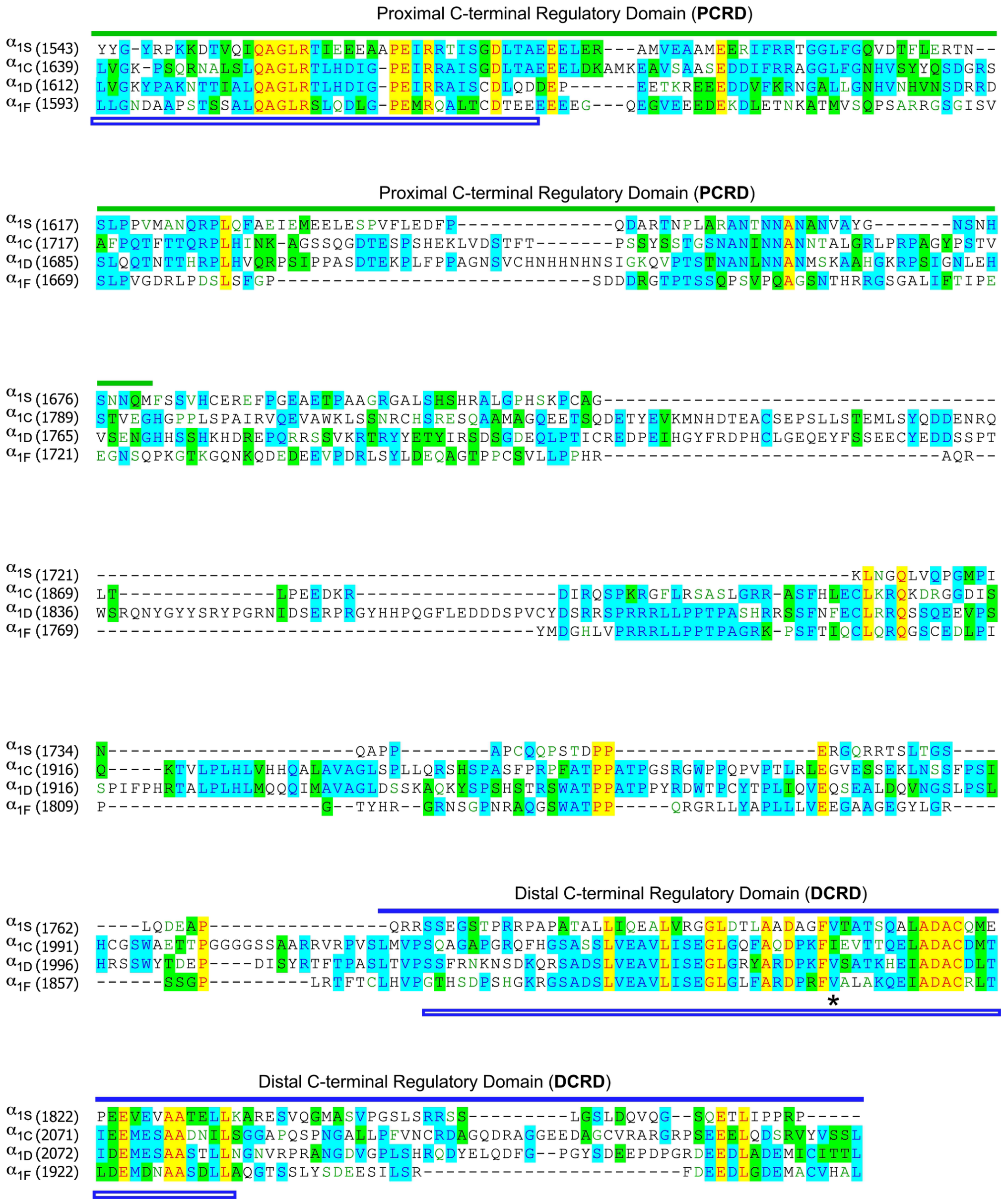


**Supplementary Figure 2S2. Alignment of the DCT domains across the Ca$_V$1 family**

The DCT sequences were aligned for Ca$_V$1.1 ($\alpha_{1S}$, NM_001101720.1), Ca$_V$1.2 ($\alpha_{1C}$, AF465484.1), Ca$_V$1.3 ($\alpha_{1D}$, EU363339.1) and Ca$_V$1.4 ($\alpha_{1F}$, NP005174), with GenBank accession numbers in parentheses. Key domains of PCRD (green) and DCRD (blue) are indicated with solid lines. The black asterisk identifies the valine residue (i.e., unveiled by Valine to Alanine or V/A mutation) that is essential for DCRD functionality. Critical topology-defining regions within the AlphaFold3-predicted DCT structure are marked by double blue lines. Sequence homology from high to low is categorized as yellow (fully identical), cyan (partially conserved), or green (similar) and – (gapped or different).

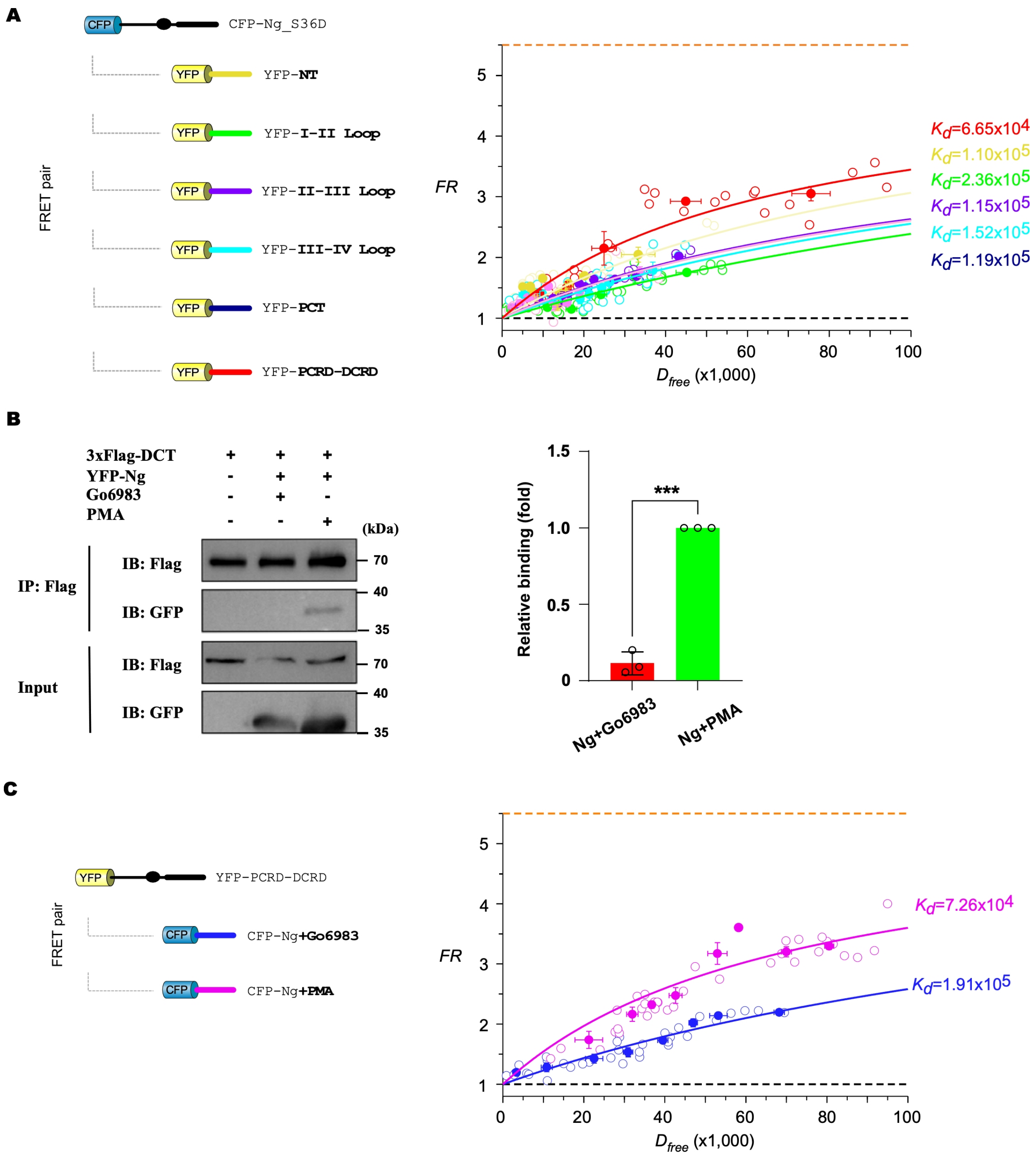


**Figure 2S3. Additional data on Ng and DCT binding**

(**A**) Binding affinities between various intracellular domains of $Ca_V1.3$ and Ng_S36D measured by 2-hybrid 3-cube FRET. The binding affinity between YFP-PCRD-DCRD (red, representing DCT) and CFP-Ng_S36D is stronger than the other FRET pairs: YFP-tagged I-II Loop, II-III Loop, III-IV Loop, or PCT, and CFP-Ng_S36D.

(**B**) Co-IP experiments for DCT (fused with the Flag tag) and Ng, treated with 0.5 μM Go6983 or 0.4 μM PMA respectively. Representative blots (left) and statistical summary (right).

(**C**) FRET 2-hybrid binding assays for CFP-Ng and YFP-PCRD-DCRD treated with 0.4 μM PMA or 0.5 μM Go6983. Each point represents the averaged values over five individual data points (adjacent to each other).

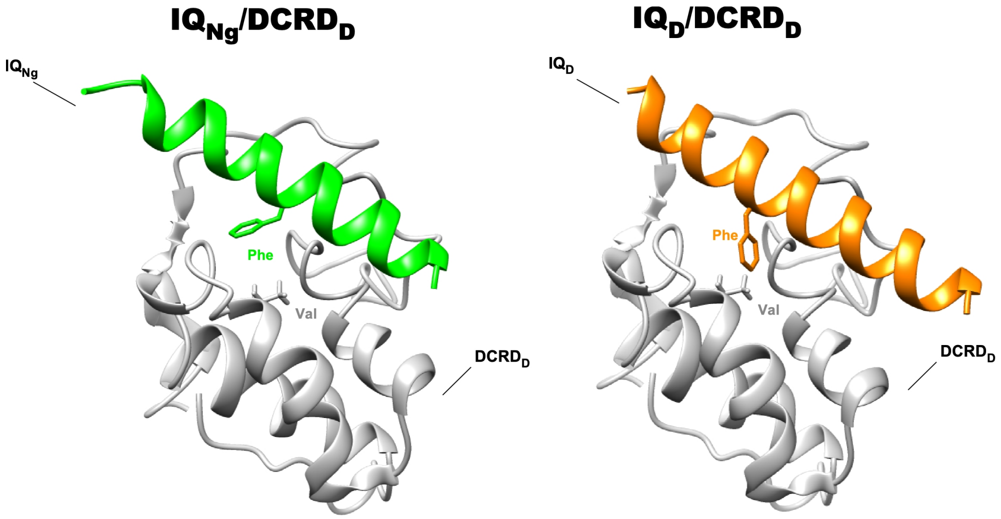


**Figure 4S1. Structural illustration of the IQ/DCT interface**

Homology structures of IQ$_{Ng}$ (green) and IQ$_D$ (orange) interfacing with Ca$_V$1.3 DCRD (gray). The side chains of Phe of IQ$_{Ng}$ or IQ$_D$ and Val of DCRD$_D$ are shown for potential hydrophobic interactions.

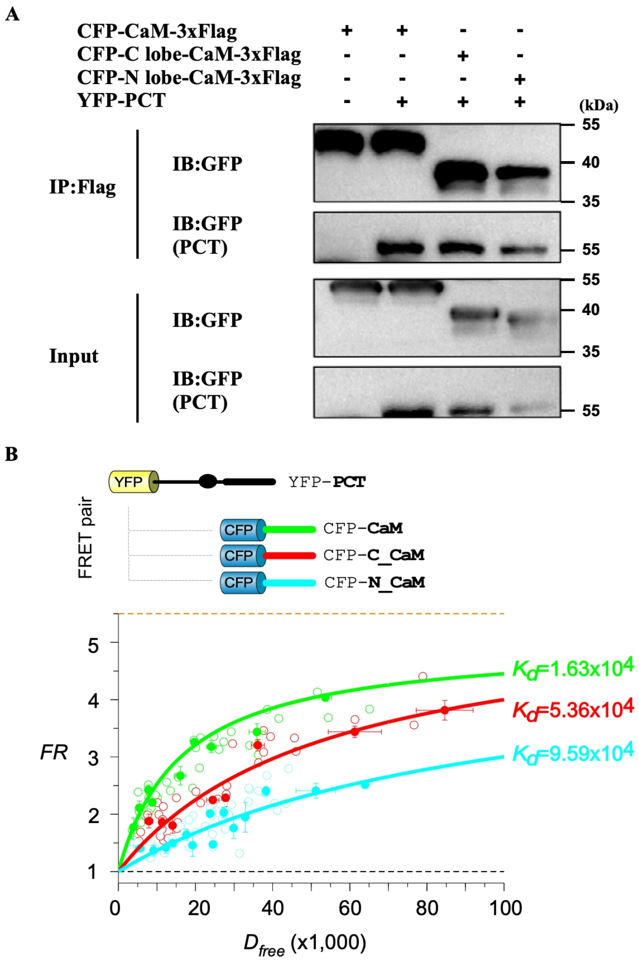


**Figure 6S1. Lobe-specific interactions between apoCaM and $Ca_V1$ PCT**

(**A**) Co-IP results regarding the interactions between YFP-PCT and CFP-CaM-3xFlag, CFP-C_CaM-3xFlag, or CFP-N_CaM-3xFlag (N-lobe of CaM). HEK293 cells were transfected with CFP-CaM, C_CaM (C-lobe of CaM) or N_CaM (N-lobe of CaM) tagged with Flag, either alone or in the presence of YFP-PCT. Cells were subjected to Co-IP 48 hours later using anti-Flag magnetic beads, followed by western blotting with anti-GFP antibody. Input lanes represent 10% of the total protein lysate.

(**B**) FRET results for the interactions between YFP-PCT and CFP-CaM, CFP-C_CaM, or CFP-N_CaM. YFP-PCT and CFP-CaM formed the binding curve of $FR$-$D_{free}$ (green) with an apparent affinity ($K_d$ = 1.63× $10^4$), compared with the binding curve for YFP-PCT and CFP-C_CaM ($K_d$ = 5.36 × $10^4$, red), and the binding curve for YFP-Ng_S36D and CFP-N_CaM ($K_d$ = 9.59 × $10^4$, cyan).

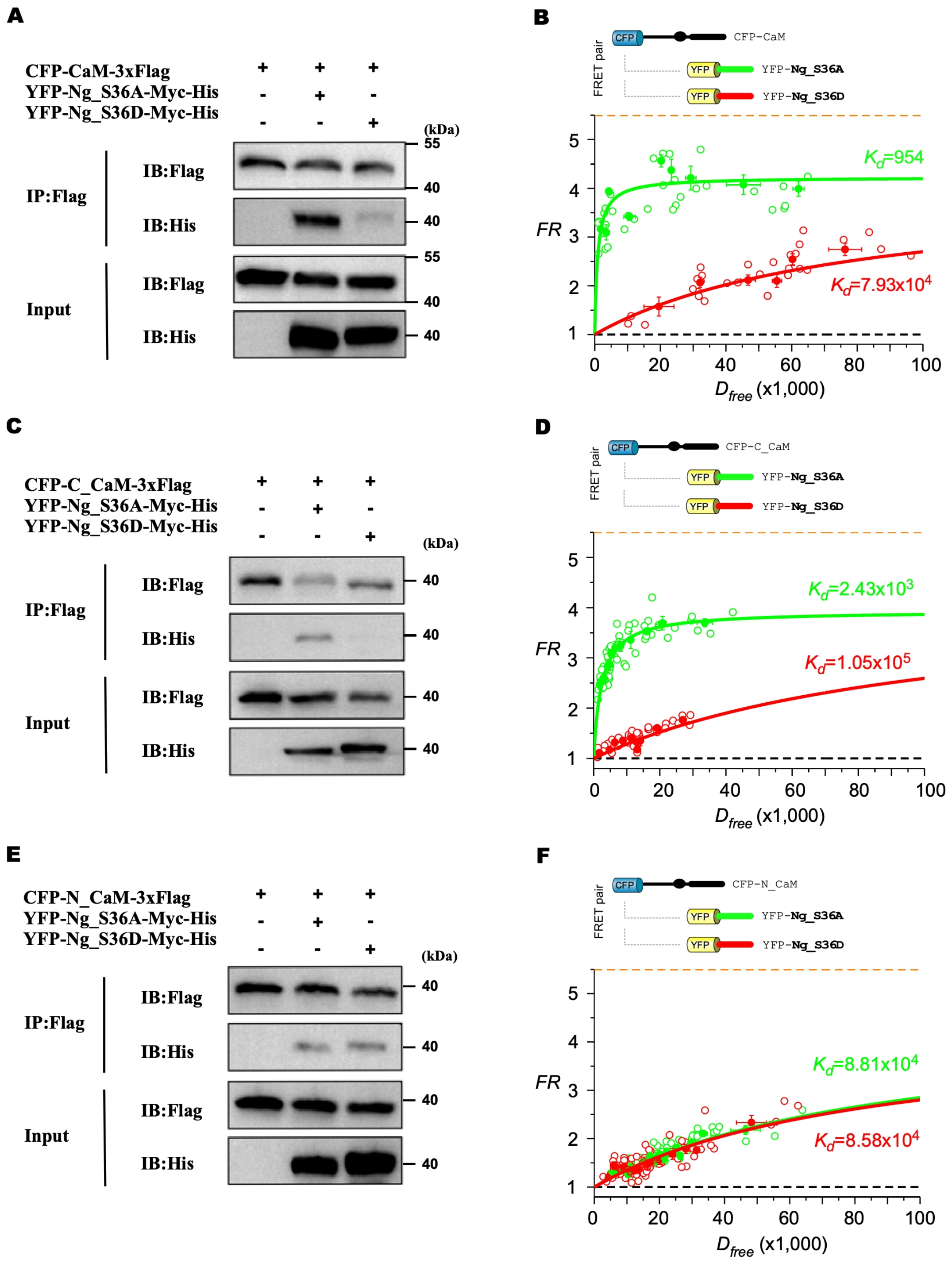


**Figure 6S2. The interactions between Ng and the two lobes of apoCaM**

(**A**) Differential interactions between unphosphorylated and phosphorylated Ng with CaM by Co-IP. HEK293 cells were transfected with CFP-CaM tagged with Flag, either alone or in the presence of YFP-Ng_S36A-Myc-His or YFP-Ng_S36D-Myc-His. Cells were subjected to Co-IP

48 hours later using anti-Flag magnetic beads, followed by western blotting with anti-His antibody or anti-Flag antibody. Input lanes represent 10% of the total protein lysate.

(**B**) Differential interactions between unphosphorylated and phosphorylated Ng with CaM by FRET. YFP-Ng_S36A and CFP-CaM formed the binding curve of $FR\text{-}D_{free}$ (green) with an apparent affinity ($K_d$ = 954); and the binding between YFP-Ng_S36D and CFP-CaM was much weaker ($K_d = 7.93 \times 10^4$, red).

(**C** and **D**) Interactions between the C-lobe of CaM (C_CaM) and pNg. Co-IP experiments were conducted between CFP-C_CaM-3xFlag, either alone or in the presence of YFP-Ng_S36A-Myc-His or YFP-Ng_S36D-Myc-His (**C**). FRET binding curves were obtained between YFP-Ng_S36A and CFP-C_CaM with an apparent affinity ($K_d = 2.43 \times 10^3$), and between YFP-Ng_S36D and CFP-C_CaM ($K_d = 1.05 \times 10^5$) (**D**).

(**E** and **F**) Interactions between the N-lobe of CaM (N_CaM) and pNg. Co-IP experiments were conducted between CFP-N_CaM-3xFlag, either alone or in the presence of YFP-Ng_S36A-Myc-His or YFP-Ng_S36D-Myc-His (**E**). FRET binding curves were obtained between YFP-Ng_S36A and CFP-N_CaM with an apparent affinity ($K_d = 8.81 \times 10^4$), and between YFP-Ng_S36D and CFP-N_CaM ($K_d = 8.58 \times 10^4$) (**F**).

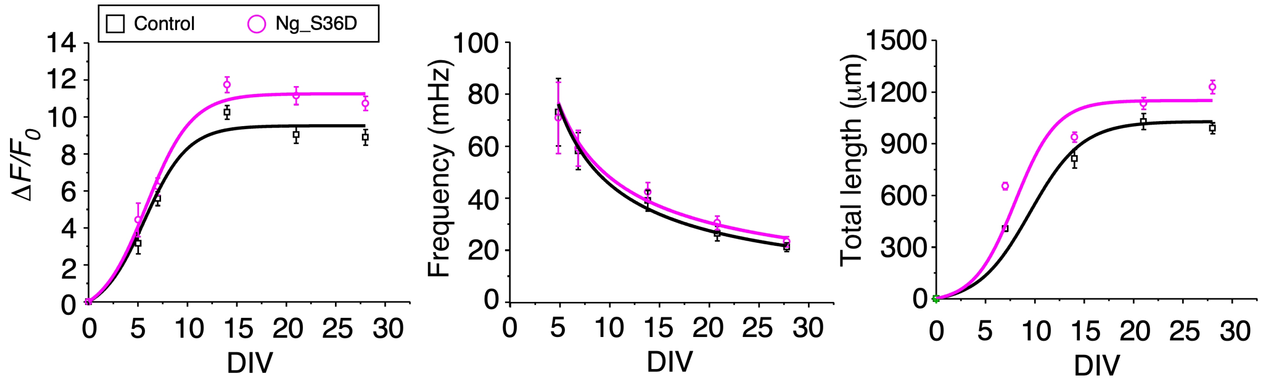


**Figure 7S1. Additional indices for the effects of overexpressing pNg on neurons**

Additional parameters related to **Figure 7C** are described in detail. These include: the time-dependent profiles of the peak amplitude ($\Delta F/F_0$), the temporal profiles of the average frequency of $Ca^{2+}$ oscillations, and the time-dependent profiles of neurite outgrowth, which are described by the correlation between the total length per single neuron (μm) and the developmental stage (DIV). All data points are presented as Mean ± SEM.

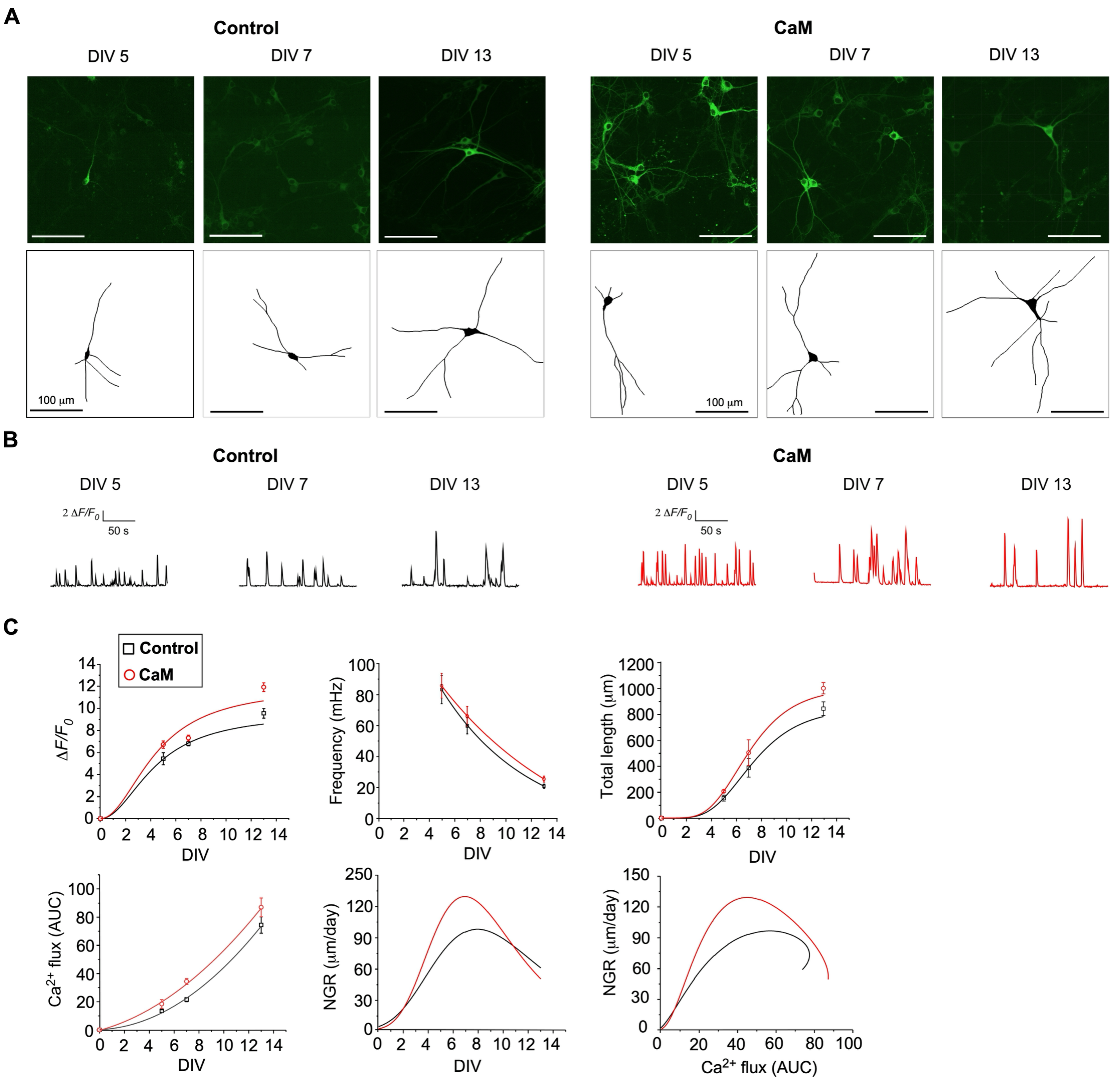


**Figure 7S2. Overexpressing CaM facilitates $Ca^{2+}$ oscillations and neurite outgrowth**

(**A** and **B**) Neurite tracing (**A**) and GCaMP-X/$Ca^{2+}$ imaging (**B**) from the control neurons versus the neurons overexpressing CaM.

(**C**) Statistical summary to compare the parameters for $Ca^{2+}$ oscillations and neurite outgrowth with (red) or without (black) CaM overexpression. Time-dependent profiles of the peak amplitude ($\Delta F/F_0$), the average frequency (in mHz), and the total neurite length per single neuron (in μm). The temporal profiles of $Ca^{2+}$ influx (AUC, $\Delta F/F_0$ per min) and neurite growth rate (NGR, μm per day) over the development stage (DIV), based on which the relationship between AUC and NGR is depicted. All data points are presented as Mean ± SEM.

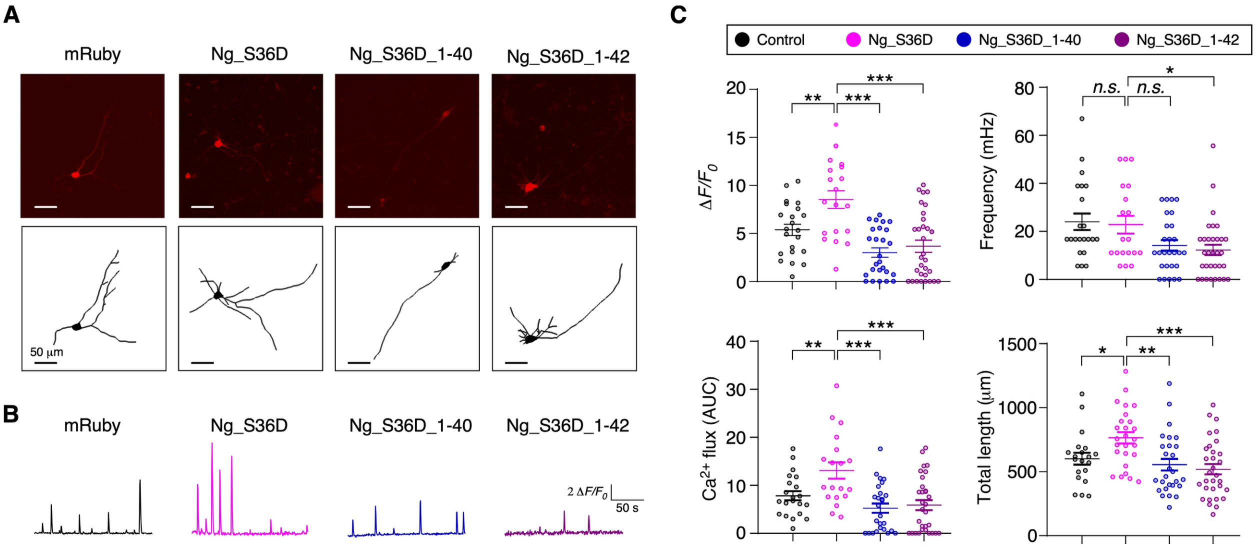


**Figure 8S1. Effects of WT and mutant pNg on hippocampal neurons**

(**A** and **B**) Neurite tracing and GCaMP imaging for the control neurons (mRuby) versus the neurons expressing Ng_S36D, Ng_S36D_1-40 or Ng_S36D_1-42.

(**C**) Key indices of $\Delta F/F_0$, the frequency, AUC and the total length were calculated and compared between the WT and mutants of Ng_S36D. All statistical data are given as Mean ± SEM. One-way ANOVA followed by Dunnett for post hoc test: *, $p < 0.05$; **, $p < 0.01$; ***, $p < 0.001$; n.s., $p > 0.05$.
